# The Age-Dependent Resident Myonuclear Multi-Omic Response to a Skeletal Muscle Hypertrophic Stimulus

**DOI:** 10.1101/2025.10.29.685384

**Authors:** Pieter J. Koopmans, Ronald G. Jones, Ana Regina Cabrera, Francielly Morena, Nicholas P. Greene, John J. McCarthy, Ahmed Ismaeel, Yuan Wen, Kevin A. Murach

## Abstract

A detailed analysis of how muscle fiber nuclei (myonuclei) respond to a hypertrophic stimulus would provide a critical step toward understanding compromised skeletal muscle plasticity with age. We used recombination-independent doxycycline-inducible myonucleus-specific fluorescent labelling, tissue RNA-sequencing, myonuclear DNA methylation analysis, multi-omic integration, and single myonucleus RNA-sequencing to define the molecular characteristics of adult (6-8 month) and aged (24 month) murine skeletal muscle after acute mechanical overload (MOV). In adult and aged MOV muscles, we found that: 1) similarities in the transcriptional response to loading – specifically in metabolism genes – were partly explained by a post-transcriptional microRNA-mediated mechanism, which we corroborated using an inducible muscle fiber-specific *miR-1* knockout model, 2) differences in age-dependent transcriptional responses were linked to the magnitude and location of differential DNA methylation in resident myonuclei, specifically around hypertrophy-associated genes such as *Myc*, *Runx1*, *Mybph*, *Ankrd1,* collagen genes, and minichromosome maintenance genes, 3) adult and aged resident myonuclear transcriptomes had differing enrichment for innervation-related transcripts as well as unique transcriptional profiles in an *Atf3+* “sarcomere assembly” population after MOV, and 4) cellular deconvolution analysis supports a role for neuromuscular junction regulation in age-specific hypertrophic adaptation. These data are a roadmap for uncovering molecular targets to enhance aged muscle adaptability.

## Introduction

Resistance exercise training is recognized as the most effective tool to induce muscle growth in humans^1–3^. Resistance training is also the most successful and accessible strategy to attenuate the natural and inevitable loss of skeletal muscle mass and function that ensues throughout the lifespan, termed age-related sarcopenia^1,4–6^. Unfortunately, the efficacy of resistance training is blunted in aged compared to younger adult human populations^7–9^. This observation is corroborated by murine models of muscle hypertrophy such as synergist ablation-induced mechanical overload (MOV) and progressive weighted wheel running (PoWeR), where aged mice have attenuated hypertrophic outcomes compared to young^10,11^. One barrier to pinpointing the molecular mechanism(s) for an age-dependent reduction of muscle plasticity is the syncytial nature of skeletal muscle fibers (myofibers). Myofibers are massive individual cells that contain hundreds to thousands of individual and often specialized nuclei (myonuclei). Beyond the contractile myofibers, skeletal muscle comprises a heterogenous cellular milieu including numerous non-muscle cell types such as immune cells, fibroadipogenic progenitors (FAPs), fusogenic muscle stem cells (satellite cells), and endothelial cells, among others. In whole tissue, the molecular profiles of these supporting cell types can disguise processes occurring specifically within myofibers, which is the cell type that adapts and physically grows in response to mechanical stimuli. Furthermore, individual myonuclei within a given muscle fiber, or within a fiber of a specific myosin heavy chain (MyHC) type (e.g. slow versus fast) may display unique impairments with aging. Manually isolating individual muscle fibers to interrogate molecular contributors to muscle plasticity is tedious and low-throughput and does not capture the complexity of individual myonuclear responses. Manual isolation of fibers may also be subject to contamination by adherent mononuclear cell types; this is especially true during times of stress characterized by mononuclear cell infiltration and a dramatic shift in muscle nuclear proportion away from myonuclei^12^. For these reasons, how myonuclei regulate muscle adaptation at the molecular level across the lifespan remains poorly understood. This lack of detailed information represents a significant gap in the skeletal muscle aging literature^13,14^.

To overcome the technical barriers of assessing muscle-fiber specific molecular profiles, we used a doxycycline-inducible genetically modified mouse model called HSA-GFP (HSA = human skeletal actin promoter; GFP = green fluorescent protein)^12,15^. This model is a recombination-independent tool that allows for fluorescent myonuclear labeling *in vivo* and sorting-based isolation of myonuclei with high specificity. Carefully timed myonuclear labeling with the HSA-GFP model eliminates the possibility of capturing nuclei from other cell types as well as myonuclei derived from newly fused satellite cells. This approach allows us to focus specifically on resident myonuclei, which have the greatest influence on initiating and driving the muscle fiber hypertrophic process^14^. Using this tool, we sought to define the resident myonuclear molecular response to a known hypertrophic stimulus (MOV) in mice according to age: 6-8-month adult versus 24-month aged. MOV shares many of the same molecular responses to acute resistance exercise in humans, and like human resistance training, has lower hypertrophic efficacy with aging^9,16–18^. We combined bulk tissue RNA-sequencing, myonucleus specific DNA methylation, multi-omic integration, and single myonucleus RNA-sequencing (smnRNA-seq) to provide a detailed portrait of how the early phase of loading-induced muscle growth is regulated by resident myonuclei (Figure 1). These data serve as a molecular resource to the skeletal muscle and aging research communities and provide fundamental insight into how muscle plasticity can be compromised late in life.

**Figure 1.**
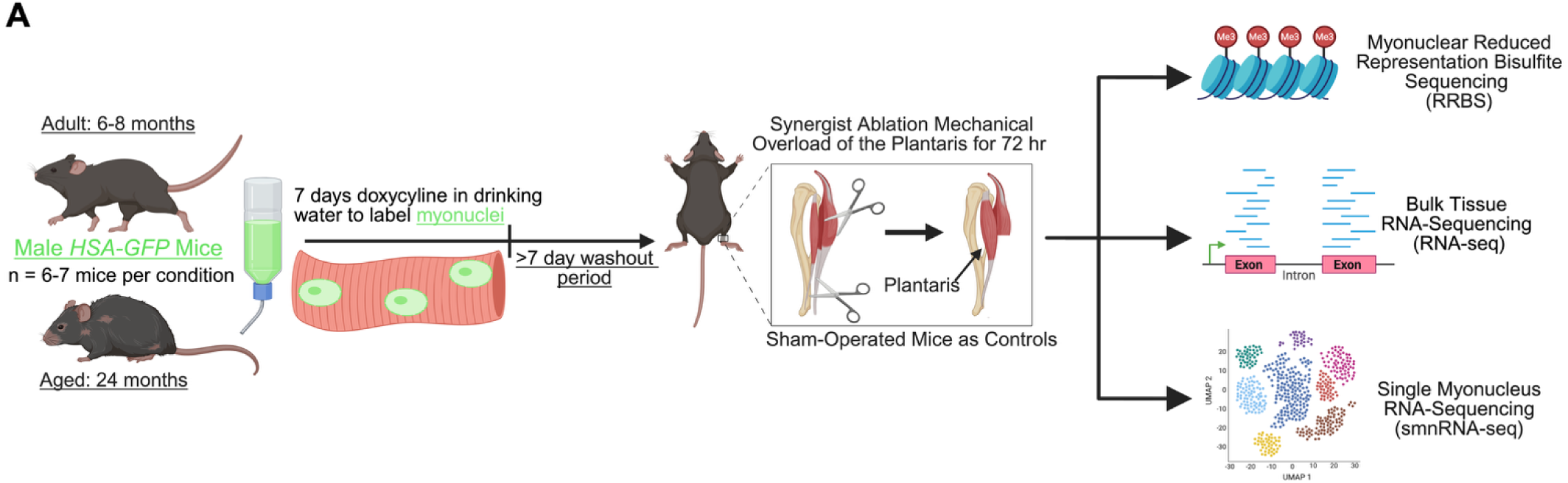
(A) Schematic overview of experimental design.

## Results

### The effect of age on the skeletal muscle transcriptome in sedentary mice

To assess baseline differences in muscle with aging, we compared whole muscle gene expression using bulk RNA-sequencing data from plantaris muscles between adult (6-8-month, n=6) and aged (24-month, n=7) male sham surgery control mice (Supplemental File 1). Principal component analysis (PCA) revealed distinct differences between the transcriptomes of adult and aged skeletal muscle (Figure 2A). Notably, there is greater dispersion of data in aged sham versus adult sham, indicative of more transcriptional variability and/or stochasticity when aged, in agreement with prior work^19^. We identified 3,200 differentially expressed protein-coding genes (DEGs) between aged sham versus adult sham mice (1,565 upregulated; 1,635 downregulated, adj. *p*<0.05, Figure 2B). Background corrected gene set enrichment analyses (GO Biological Processes) were performed separately on up- and down-regulated genes^20,21^. With age, upregulated GO terms were primarily associated with mitochondrial gene expression, translation, and ribosome structural constituent genes (Figure 2E). These pathways were defined by numerous ribosomal protein genes (*Rps and Rpl)* and eukaryotic initiation factor (*Eif)* subunits. Although seemingly counterintuitive, our observation of upregulated ribosomal regulation in aged mouse muscle is consistent with prior work showing ribosomal RNA and proteins are elevated in aged human muscle at rest^22^. Furthermore, it is becoming clear that hyperactive mTORC1 signaling and elevated protein synthesis with aging is a driver of impaired proteostasis and sarcopenia, and that partial inhibition of mTORC1 with rapamycin reverses age-related muscle deficits in sedentary animals^23–27^. Downregulated GO terms were related to skeletal system development, response to growth factors, and angiogenesis, but were dominated by terms related to extracellular matrix (ECM) organization (Figure 2F). Downregulation of these gene classes with aging may signal a loss of cellular identity, accompanied by impaired ECM remodeling that is known to occur in aging skeletal muscle^28–32^.

**Figure 2.**
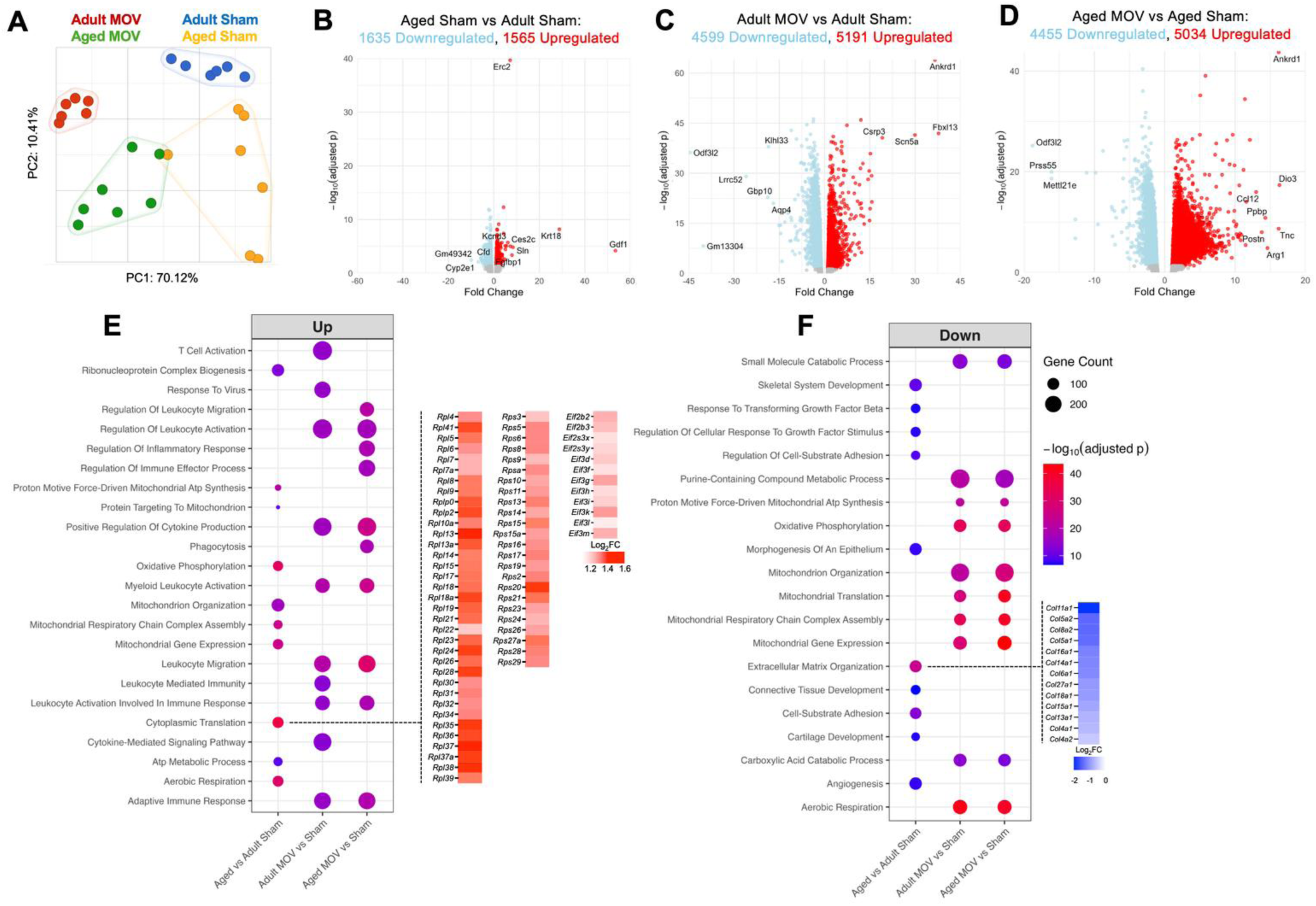
(A) Principal component analysis of top 2000 features across experimental conditions. Volcano plots of (B) Aged Sham vs Adult Sham, (C) Adult MOV vs Adult Sham, (D) Aged MOV vs Aged Sham (adj. *p*-value < 0.05). (E) Plot showing top 10 GO biological processes pathways based on up-regulated genes from each comparison. (F) Plot showing top 10 GO biological processes pathways based on down-regulated genes from each comparison. MOV = mechanical overload. FC = fold- change.

### Conservation of immune-oriented and metabolic transcriptional profiles between adult and aged mouse muscle in response to an acute hypertrophic stimulus

We next sought to characterize the transcriptome of skeletal muscle (plantaris) in response to acute (72 hour) synergist ablation-induced MOV in adult and aged male mice (Supplemental File 1). This early timepoint precedes *bona fide* muscle fiber growth but is critical for the long-term muscle growth response^14,33^. PCA revealed greater variability in aged animals in response to MOV (Figure 2A). In adult mice, 9,700 DEGs were identified in MOV versus sham (5191 upregulated, 4599 downregulated, adj. *p*<0.05, Figure 2C). In aged mice, 9,489 DEGs were identified in MOV versus sham (5,034 upregulated, 4,455 downregulated, adj. *p*<0.05, Figure 2D). GO pathway analyses (top 10 biological processes shown) from DEGs in adult and aged MOV conditions, respectively, showed generally similar enriched GO terms. Upregulated GO pathways in adult and aged were overwhelmingly related to immune responses (Figure 2E). Induction of an immune response at the tissue level in muscle is unsurprising given pronounced inflammatory cell appearance at this early stage of MOV^12^. Downregulated GO terms were dominated by metabolic-related processes, including mitochondrial organization, cellular respiration, and energy metabolism (Figure 2F). The signature of downregulated mitochondrial-related genes may be a sign of the Warburg effect (or aerobic glycolysis) similar to what is observed in cancer cells during rapid growth^34,35^. We previously demonstrated this effect to occur during rapid MOV-induced muscle hypertrophy of the plantaris muscle in adult mice^36^. This response results in a biasing of substrates towards “aerobic glycolysis” and the pentose phosphate pathway to support biomass accumulation (nucleotide synthesis) that occurs concomitant with reduced mitochondrial respiration^36,37^. This Warburg-like metabolic signature appears to be conserved with MOV across ages during MOV.

### Regulation of Golgi-related and mitochondrial gene expression by *miR-1* independent of age

Our recent work suggests that miRNAs are powerful regulators of the skeletal muscle transcriptome and the metabolic response to MOV^36,37^ as well as exercise training responses in murine skeletal muscle^38^. Given our recent findings, we used DIANA-TarBase, an experimentally validated miRNA:gene expression database^32^, to guide our analysis of influential miRNAs affecting gene expression with MOV independent of age. The microRNA predicted to be most explanatory for gene upregulation with MOV regardless of age was the myomiR *miR-1* (Figure 3A). *miR-1* is a muscle-enriched microRNA hypothesized to act as a “molecular brake” on muscle growth given its downregulation coincides with hypertrophic stimuli in murine and human muscle^37,38,40–42^. In the present study, *miR-1* levels were precipitously lower with MOV regardless of age (Figure 3B). To determine how *miR-1* influences muscle gene expression in the plantaris muscle, we used a skeletal muscle-specific Cre-mediated tamoxifen-inducible *miR-1* knockout (KO) mouse model, HSA-miR-1 (Figure 3C, Supplemental Figure 1A, Supplemental File 2)^40^. In middle-aged experimental mice (12-14 months at euthanasia), *miR-1* was depleted in the plantaris by >95% relative to tamoxifen-treated controls (Figure 3D). RNA-seq revealed 875 upregulated and 756 downregulated genes after *miR-1* depletion (Figure 3E and 3F). Gene set enrichment analyses indicated upregulation of Golgi membrane and endomembrane-related genes (Figure 3E) with *miR-1* depletion. This expression profile aligns with a previous report on *miR-1* overexpression in skeletal muscle during MOV, where *miR-1* induction blunted hypertrophy (Figure 3E and 3G)^42^. In our *miR-1* KO RNA-seq, ADP-ribosylation factor 4 (*Arf4),* which regulates endosome recycling and intracellular trafficking^43,44^, is a validated AGO2 eCLIP-seq target of miR-1 in human muscle tissue^40^, as are cell-cycle regulators that are implicated in tumorigenesis: *Azin1* and *Ptpn1*^45–47^ (Supplemental Figure 1). Genes related to cellular respiration and oxidoreductase activity were strongly downregulated in *miR-1* KO relative to controls (Figure 3F and 3H). There was appreciable overlap in the transcriptomes between miR-1 KO and MOV, specifically related to downregulation of mitochondrial enzymes and mitoribosome-related genes (Supplemental Figure 1B). *miR-1* has previously been shown to control glycolytic flux and mitochondrial respiration in the oxidative soleus muscle of adult mice^40^. Altogether, our data indicate *miR-1* is likely responsible for controlling the metabolic response as well as Golgi membrane remodeling processes during MOV in the fast-twitch plantaris muscle irrespective of age. A *miR-1* mediated post-transcriptionally controlled metabolic switch may help to create a permissive environment for muscle growth^42^.

**Figure 3.**
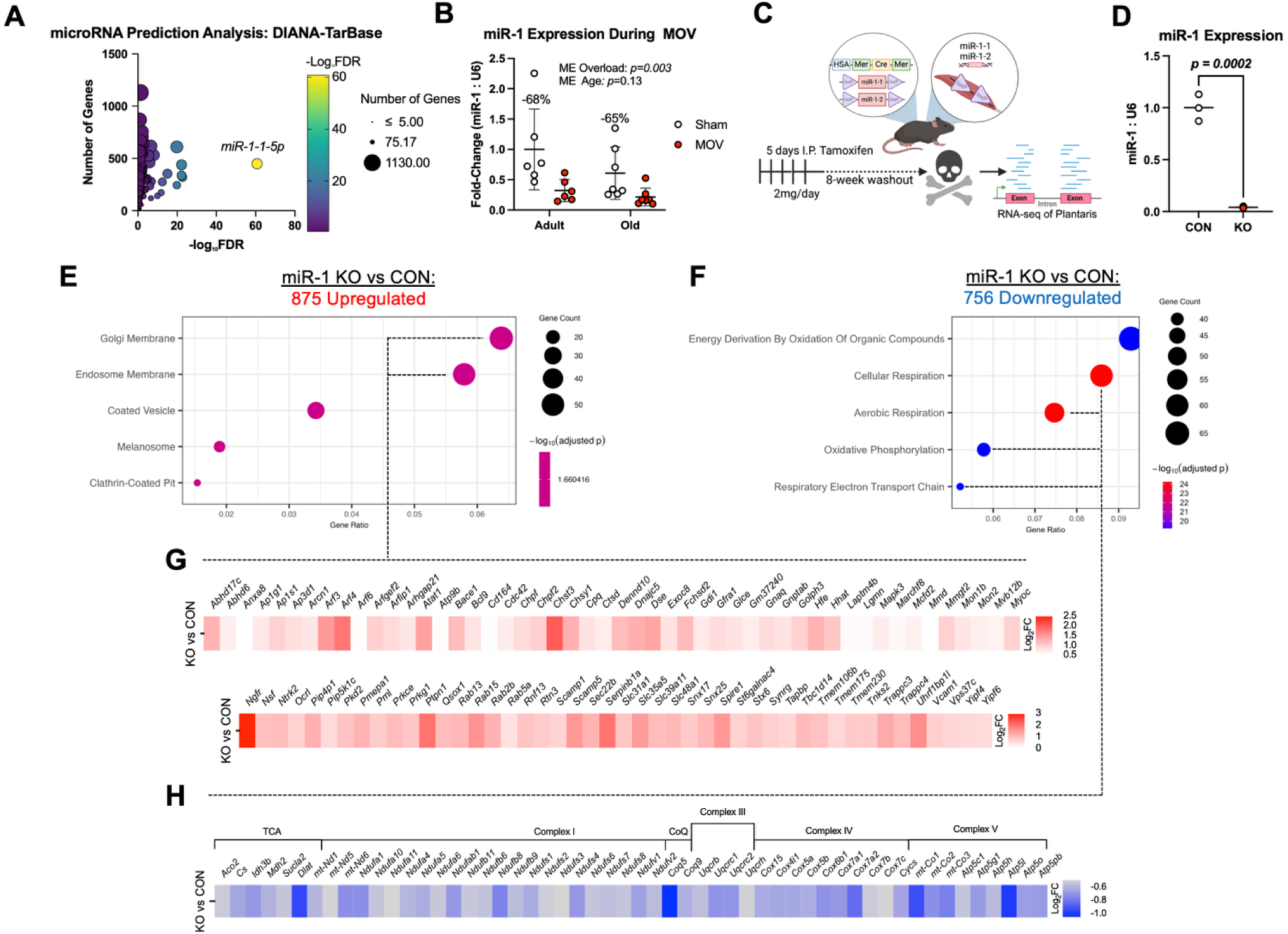
(A) DIANA-TarBase microRNA regulation prediction analysis from upregulated genes in response to MOV in both age conditions. (B) RT-qPCR of *myomiR-1* expression following MOV in adult and aged mice. (C) Experimental design summary of tamoxifen inducible muscle-specific *miR-1* knockout model (HSA-miR-1) and RNA-sequencing analysis. (D) RT-qPCR of *miR-1* expression with tamoxifen treatment in HSA-miR-1 mouse model. (E) GO Cellular Components from genes upregulated by *miR-1* KO. (F) GO Biological Processes from genes downregulated by *miR-1* KO. (G) Heatmap of upregulated endomembrane and Golgi membrane system-related genes. (H) Heatmap of downregulated TCA cycle and ETC-related genes, MOV = Synergist-Ablation Induced Mechanical Overload. ME = main effect. CON = control. KO = knock out. TCA = tricarboxylic acid cycle. CoQ = Coenzyme Q.

### Computational deconvolution of cell type contributions to MOV in adult versus aged muscle reveals a role for the neuromuscular junction

We previously performed deconvolution analysis of bulk RNA-sequencing data using a reference single cell RNA-seq (scRNA-seq) muscle regeneration dataset to infer which cell types contribute to the global muscle MOV transcriptome in young mice (∼2 months old)^48,49^. To determine if predicted changes in cell proportion during MOV is influenced by age, we performed cellular deconvolution using scRNA-seq data from a 4-day plantaris tenotomy dataset (Figure 4A)^50^. In both adult and aged MOV, myonuclei were the major source of transcription with fibro-adipogenic progenitors (FAPs) and monocytes being the next largest components. Statistical analyses of predicted cellular proportions suggested greater abundance of glial cell and muscle satellite cell populations in adult versus aged mice during MOV (Figure 4B). In aged muscle, satellite cell activation and proliferation is impaired during muscle hypertrophy, so the present result is not unexpected^51–53^. Glial cells promote neuromuscular junction integrity during denervation and express genes implicated in the ECM^54^. Glial cells may also coordinate with satellite cells to support neuromuscular junction repair during muscle injury^54^. Recent evidence suggests subsets of aged satellite cells are enriched for neuromuscular genes during MOV, potentially contributing to innervation^55^. We have previously shown how depletion of satellite cells in adult muscle results in excessive collagen accumulation concomitant with blunted long-term muscle growth^33,56,57^. Collectively, our analysis suggests satellite cells, glial cell, and innervation-related processes may be compromised during MOV with aging, which could in part explain age-dependent muscle plasticity.

**Figure 4.**
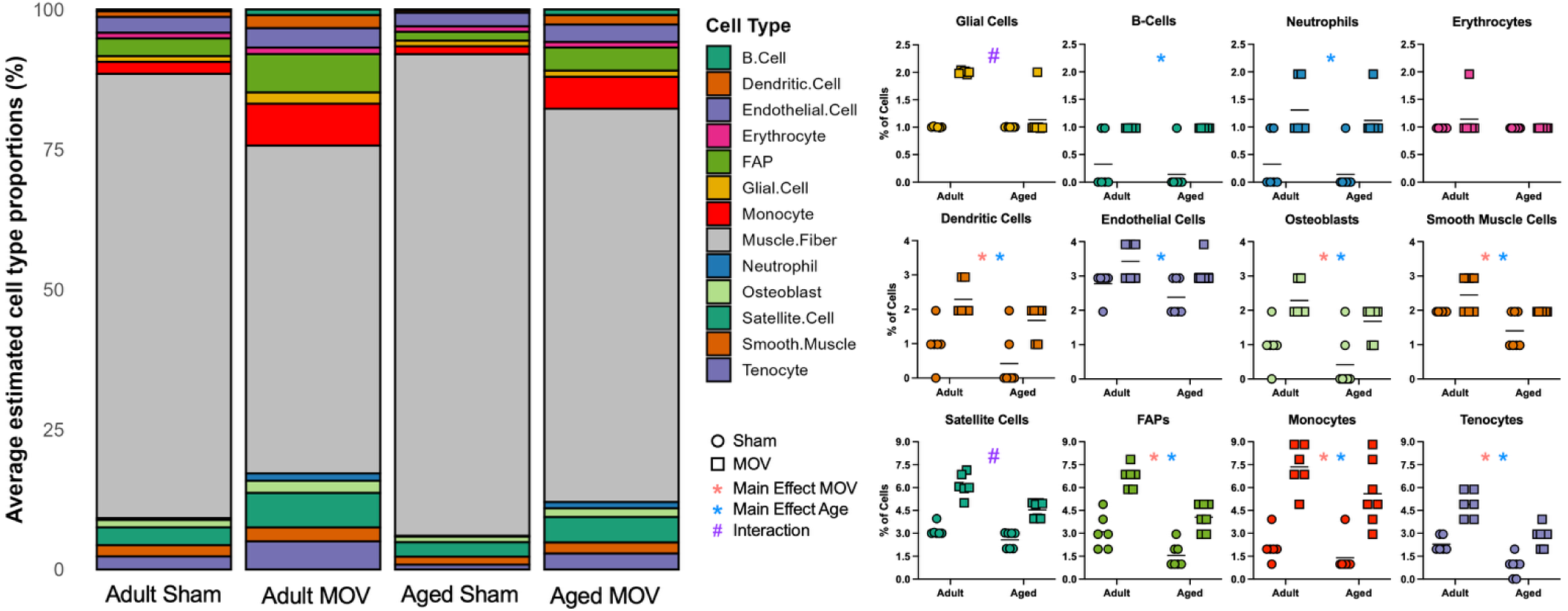
(A) Digital deconvolution of cell proportion from MOV bulk RNA-sequencing data using Granulator and Zhang et al., *Cell Reports*, 2024 as a reference scRNA-seq dataset to delineate contributions to the global transcriptome. (B) Predicted proportion of cell types. Main effect of MOV = red asterisk. Main effect of Age = blue asterisk. Interaction = #.

### Distinctions in age-related gene expression during MOV

Broad transcriptional patterns at the tissue level were generally conserved between adult and aged mice with MOV (Figure 2, Figure 5A); however, there were still numerous noteworthy distinctions. The overall transcriptional response to acute MOV was profound regardless of age, so despite significant overlap in DEGs (>8,300 shared between adult and aged MOV), there remained >1,400 DEGs exclusive to adult mice and >1,100 DEGs exclusive to aged mice (adj. *p*<0.05 relative to respective controls, Figure 5A). These genes may be important for explaining compromised muscle plasticity when aged. Twenty-three genes were oppositely regulated in adult versus aged mice after MOV (adj. *p*<0.05). Figure 5B shows genes that were upregulated in adult and downregulated in aged, or vice versa. One such gene is *Casp12*, which was downregulated in adult and upregulated in aged, is an initiator of caspase leading to apoptosis. Deletion of *Casp12* preserves muscle function and reduces signs of muscle degeneration in *mdx* mice^58^. *Sema3a* was also downregulated with MOV in adult but upregulated in aged MOV. *Sema3* is implicated in muscle mass regulation, where overexpression in IIB muscle fibers - the predominant myosin heavy chain isoform in plantaris muscle - prevents fusion of *Tw2+* myogenic progenitor cells (MPCs) to the muscle fiber^59^. Genetic ablation of *Tw2+* MPCs causes IIB fiber atrophy^60^. Perhaps the upregulation of *Sema3a* in aged mice during MOV prevents fusion of this IIB-specific myogenic progenitor population and affects muscle adaptation. Importantly, *Twist2*+ cells are also found in human skeletal muscle and are responsive to aging and resistance exercise^61,62^. Enriched GO biological process terms from upregulated DEGs exclusive to adult MOV were related to ribonucleoprotein complex biogenesis, ribosome biogenesis, and translation at the pre-synapse (Figure 5C). Enriched GO terms from downregulated DEGs exclusive to adult MOV (Figure 5D) were related to the proteasome and endopeptidase complex. By contrast, enriched GO terms from upregulated DEGs exclusive to aged MOV mice were related to coagulation, hemostasis, and negative regulation of cell motility. There were no enriched GO terms for downregulated DEGs exclusive to aged MOV. Unique upregulated genes in aged MOV collectively point to wound repair processes (Figure 5E) and suggest an altered early response to MOV when aged.

**Figure 5.**
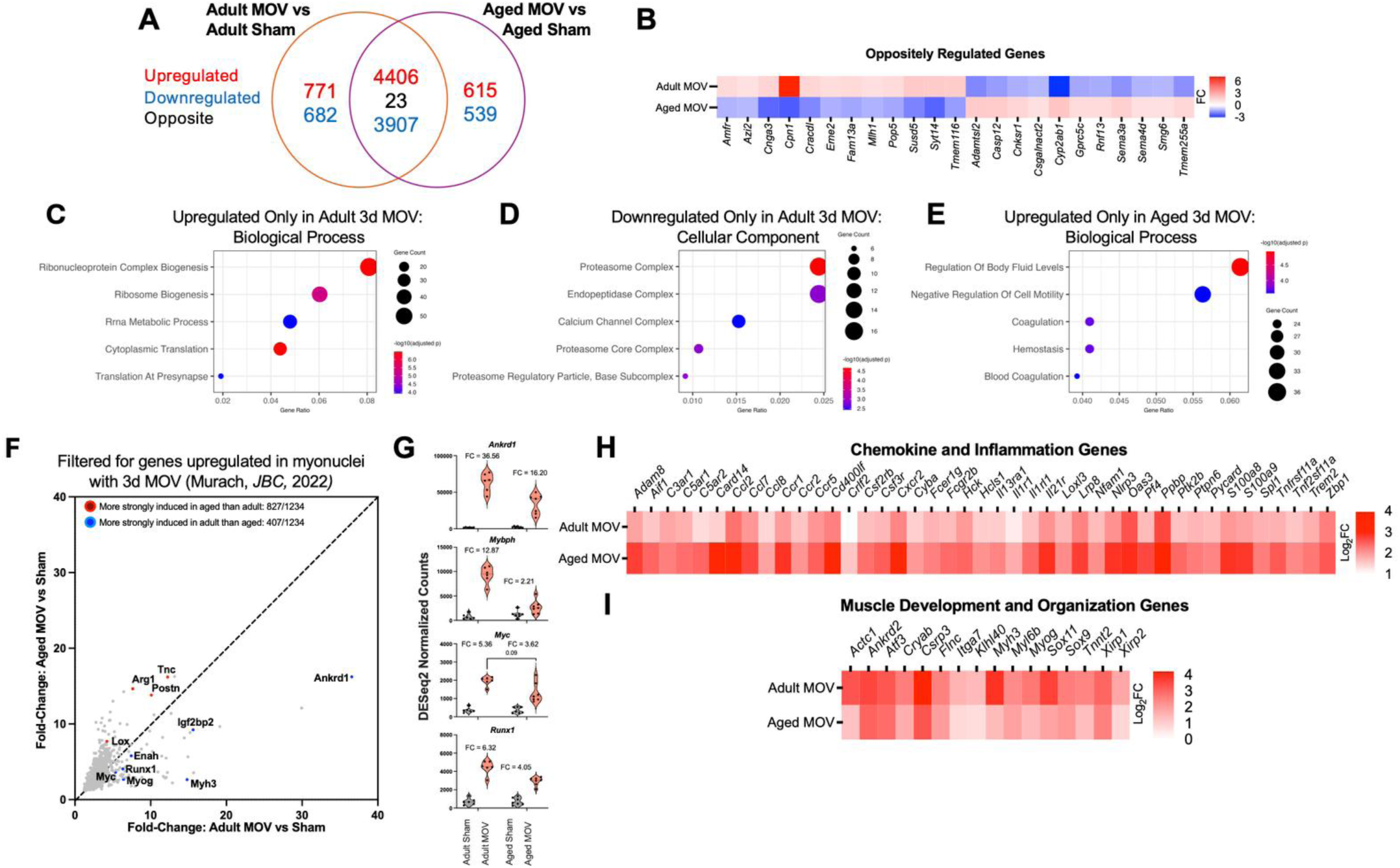
(A) Venn diagram comparing overlap in differentially expressed genes in adult and aged conditions in response to MOV. (B) Heatmap with fold changes of genes that are oppositely regulated, upregulated in adult MOV but downregulated in aged MOV, downregulated in adult MOV but upregulated in aged MOV. (C) Top 5 enriched GO Biological Processes from upregulated DEGs exclusive to adult MOV, (D) Top 5 enriched GO Biological Processes from downregulated DEGs exclusive to adult MOV. (E) Top 5 enriched GO Biological Processes from upregulated DEGs exclusive to aged MOV. (F) Fold-change induction of genes filtered for expression in myonuclei (Murach et al. *Journal of Biological Chemistry,* 2022), (G) Violin plots showing normalized counts and fold changes of select myonuclei enriched genes (Aged MOV vs Adult MOV adj. *p*-values: *Ankrd1*: <0.01; *Mybph*: <0.0001; *Myc*: 0.09; *Runx1*: 0.12), (H) Heatmap of Chemokine and Inflammation Genes. (I) Heatmap of Muscle Development and Organization-Related Genes. FC = fold-change

To further refine our examination of the response to MOV, we focused on genes previously identified to be enriched specifically in bulk myonuclei and upregulated in response to 72-hour MOV in young mice (1694 genes)^48^. There were 1,234 overlapping with the tissue RNA-seq herein, of which 827 were more strongly induced in aged muscle than adult, and 407 more strongly induced in adult muscle than aged (Figure 5F). *Arg1, Lox, Tnc,* and *Postn,* and *Timp1* were more strongly induced in aged muscle than adult (Figure 5F). *Arg1* is highly upregulated in macrophages after 4 days of MOV^63^, and *Timp1* is a macrophage-derived pro-inflammatory cytokine^64–66^. Several genes we previously identified to be enriched in myonuclei and associated with muscle mass regulation were also more strongly induced in adult than in aged mice, including *Myc, Runx1, Mybph, Ankrd1* (Figure 5G). We recently identified skeletal muscle-specific induction of *Myc,* a highly exercise-responsive Yamanaka Factor, to be sufficient for muscle growth^67^. *Myc* was previously shown to be less responsive to a muscle growth stimulus with age across species and conditions^68–71^; perhaps this age-associated attenuation contributes to impaired muscle growth^48,68,72,73^ and also explains aging-associated declines in loading-mediated ribosome biogenesis^9^. *Runx1,* while induced strongly during MOV, is similarly upregulated during muscle denervation. *Runx1* may act as a pro-hypertrophy signal as well as anti-atrophy countermeasure given depletion of *Runx1* decreases ribosome biogenesis in hematopoietic stem cells and augments muscle wasting and myofibrillar disorganization during denervation^74–76^. Chemokine and inflammation-oriented genes were more strongly induced in aged mice with MOV relative to adult (Figure 5H), while genes related to muscle development and organization were more strongly induced in adult muscle (Figure 5I). The stronger induction of muscle development and organization genes at the tissue-level with MOV when younger could be related to a more robust satellite cell response versus old. We recently provided evidence for the presence of satellite cells reinforcing muscle identity genes during lifelong wheel running^77^. Overall, distinctions in the bulk transcriptome of adult versus aged mouse muscle could contribute to reduced plasticity with loading when aged.

### The myonuclear DNA methylome is significantly altered by age and is dramatically less responsive to MOV in aged than adult muscle

Using reduced representation bisulfite sequencing (RRBS), we previously reported the myonuclear DNA methylome is altered by acute MOV in young female mice (∼2 months old)^12,36^. However, to date, the myonuclear DNA methylome response to MOV has not been defined in adult or aged muscle. We performed low-input RRBS on myonuclei isolated via fluorescence activated nuclear sorting (FANS) from the plantaris muscle of sham control and MOV mice. We initially focused on promoter CpG methylation as this is canonically linked to gene expression regulation when compared with other genomic contexts such as exons/introns^78,79^. Examining the effect of age (aged sham versus adult sham), 17,494 promoter CpG sites were differentially methylated (DM, Figure 6A&A’, adj. *p*<0.05 and >10% methylation difference, 2202 unique genes). Around 22% of age-related DM genes were both hypo- and hypermethylated (“mixed”), with the other 78% of genes being exclusively hypo- or hypermethylated. Many DM CpG sites were altered by up to ±25-50% (Figure 6A”), indicating the myonuclear DNA methylome is dramatically altered by age alone, consistent with reports in tissue of mice and humans^80–83^.

**Figure 6.**
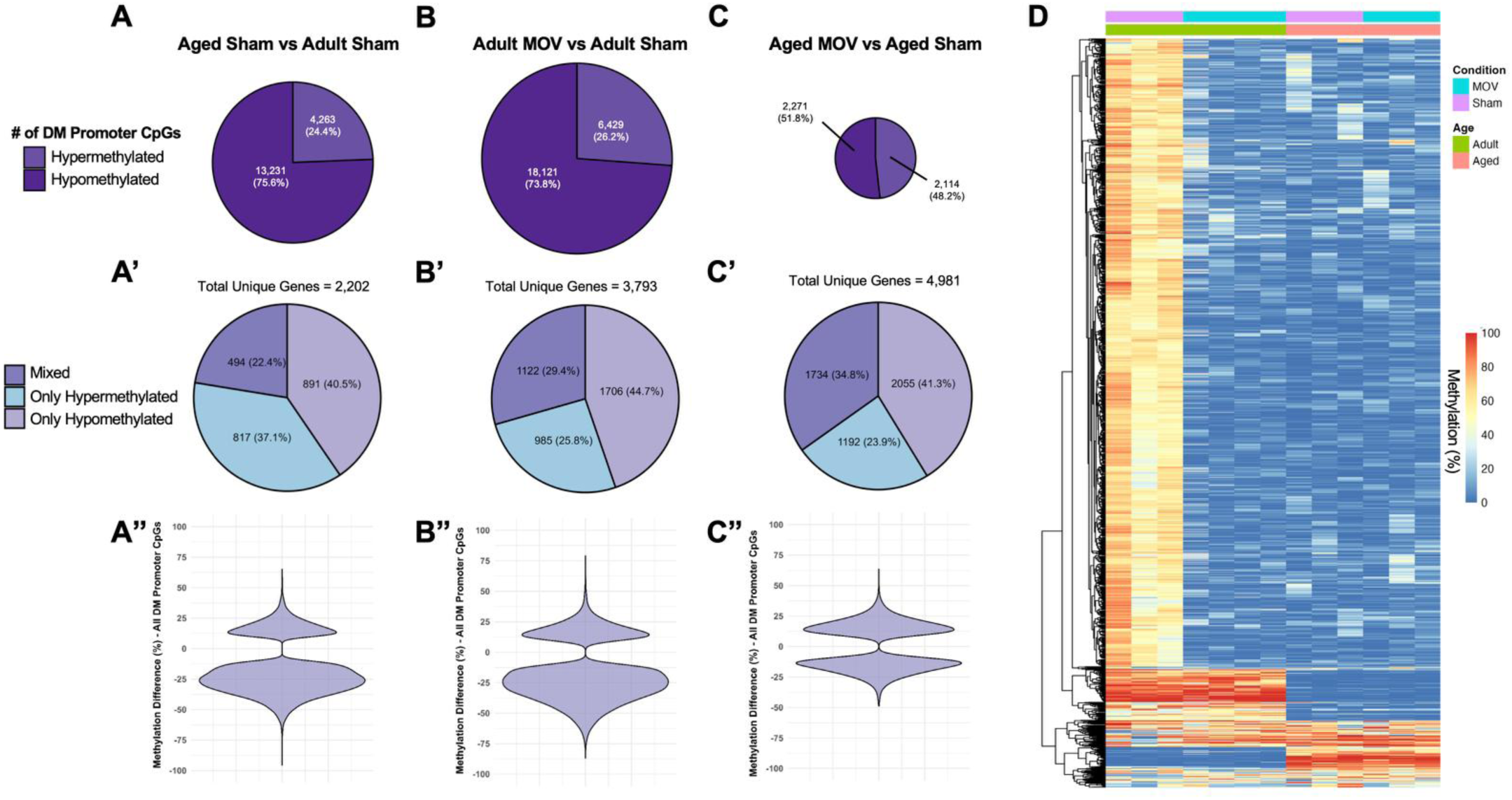
Number of promoter CpG sites with differential methylation for: (A) aged sham vs adult sham, (B) adult MOV vs adult sham, (C) aged MOV vs aged sham. Number of unique genes which are of mixed methylation status, only hypomethylated, or only hypermethylated for: (A’) aged sham vs adult sham, (B’) adult MOV vs adult sham, (C’) aged MOV vs aged sham. Violin plots of % differential methylation for: (A”) aged sham vs adult sham, (B”) adult MOV vs adult sham, (C”) aged MOV vs aged sham. (D) Hierarchical clustering heatmap of top 2000 variable DM CpG sites. DM = differentially methylated.

In adult MOV, there were 24,550 DM promoter CpGs relative to age-matched sham, whereas in aged MOV there were only 4,385 DM CpGs (Figures 6B and 6C). This striking difference suggests the myonuclear methylome is less flexible after short-term MOV in aged myonuclei, perhaps due in part to the significant alterations with age observed in resting conditions (see above). For MOV comparisons, the DM CpGs corresponded to 3,793 unique genes in adult and 4,981 unique genes in aged (Figure 6B’ and 6C’). Aged muscle had fewer differentially methylated sites but in more genes with MOV, suggesting a more uncoordinated epigenetic response to loading. Less epigenetic flexibility in aged muscle is further supported by a relatively lower percent change in methylation of individual CpG sites compared to adult (Figure 6B” and 6C”). In aged MOV, most CpG sites exhibited ±10-25% differential methylation relative to control myonuclei with few outside of that range (maximum of ±50%). In contrast, adult MOV had more CpG sites altered by ≥25% and up to ±80% methylation difference. The marked differences in the magnitude of DNA methylation responses to MOV between adult and aged mice are illustrated by the heatmap of the top 2000 most variable CpG sites in Figure 6D. It is worth noting that with MOV in adult and aged, 65-71% of genes had differential methylation in promoter regions that were only hypo- or hypermethylated, whereas 29-35% of genes had promoter CpG sites that were both hypo- and hypermethylated, or “mixed” methylation (Figure 6B’ and 6C’).

### Methylome-transcriptome integration shows epigenetic control of the muscle transcriptome with aging

Interpreting the impact of DNA methylation on gene expression can be complicated since many genes have regulatory regions with CpG sites that are of mixed methylation status (as shown in Figure 6). In an effort to decode this ambiguity, we used Binding and Expression Target Analysis (BETA), a multi-omic integration tool that uses several genomic parameters that are relevant to inferring epigenetic control of transcription^84^. BETA was first adapted by us from ChIP-seq analysis to understand how myonuclear DNA methylation regulates the transcriptome during acute MOV in young female mice^34^. In this prior work, BETA inferred the epigenetic regulation of metabolic adaptations in muscle during MOV, which we corroborated using high-resolution respirometry. We have since used this approach to understand how DNA methylation regulates the transcriptome throughout recovery from acute resistance exercise in humans^70^, transcriptional remodeling after muscle injury when aged^85^, as well as how the methylome may influence the proteome with late-life exercise training in mice^86^. BETA generates regulatory scores for overall up- or down-regulation and for individual genes, accounting for transcription start sites (TSS) proximity and magnitude of differential methylation. In some cases, the overall BETA regulation score for up- or downregulation may not be statistically significant but there can still be substantial regulation on a gene-by-gene basis, which we considered in our analysis. By combining our RRBS and transcriptome datasets and focusing first on the effect of age (6-8 month versus 24-month sham mice), BETA inferred differential expression of 1,950 genes due to myonuclear DNA methylation (Supplemental Figure 2A, Supplemental File 3). Altered pathways of DNA methylation-controlled genes largely reflected that of bulk RNA-seq (Supplemental Figure 2B and 2C), supporting the hypothesis that changes to the DNA methylome have significant regulatory influence on the muscle transcriptome as age progresses^80,82,83,87–89^.

### Methylome-transcriptome integration suggests greater epigenetic control of muscle growth-oriented genes in adult versus aged MOV

BETA predicted an overall similar number of genes to be regulated by DNA methylation in aged and adult mice in response to acute MOV (Figure 7A and 7B); however, the proportion of genes common to MOV in adult versus aged was only ∼36%, (Figure 7C, Supplemental File 3). Among common BETA target genes sharing the same overall pattern of expression (upregulated or downregulated in both ages), adult MOV featured more predicted regulatory CpG sites on average per gene, and the altered CpGs were relatively closer to the TSS compared to aged (Figure 7D). For example, in the BETA analysis, *Myc* in adult MOV had 10 predicted regulatory myonuclear CpG sites that were on average 826 base pairs (bp) from the TSS, but in aged MOV there were only 5 regulatory CpG sites that were on average 58,175 bp from the TSS (Supplemental Figure 3A). Independent from BETA integration, myonuclear RRBS data alone corroborates epigenetic effects on *Myc* with 11 significantly hypomethylated CpGs in adult MOV but only a single hypomethylated CpG site for aged MOV (Supplemental Figure 3B). This pattern holds for several other genes that had blunted transcriptomic responses to MOV when aged (see Figure 5G): *Myc, Runx1, Mybph,* and *Ankrd1* (Supplemental Figure 3C). *Myc* expression is seemingly controlled by DNA methylation in other tissues and contexts^90–94^, so differences in myonuclear DNA methylation around the *Myc* gene and others may explain blunted gene expression responses to MOV in aged versus adult. In general, a majority of common BETA targets exhibited greater fold-changes in gene expression after MOV in adult versus aged (Figure 7E). GO pathway analyses of common BETA regulated genes in adult and aged points to upregulation of actin filament organization, signal transduction, and organelle organization (Figure 7H, left) and downregulation of mitochondrial aerobic respiration and oxidative phosphorylation (Figure 7H, right).

**Figure 7.**
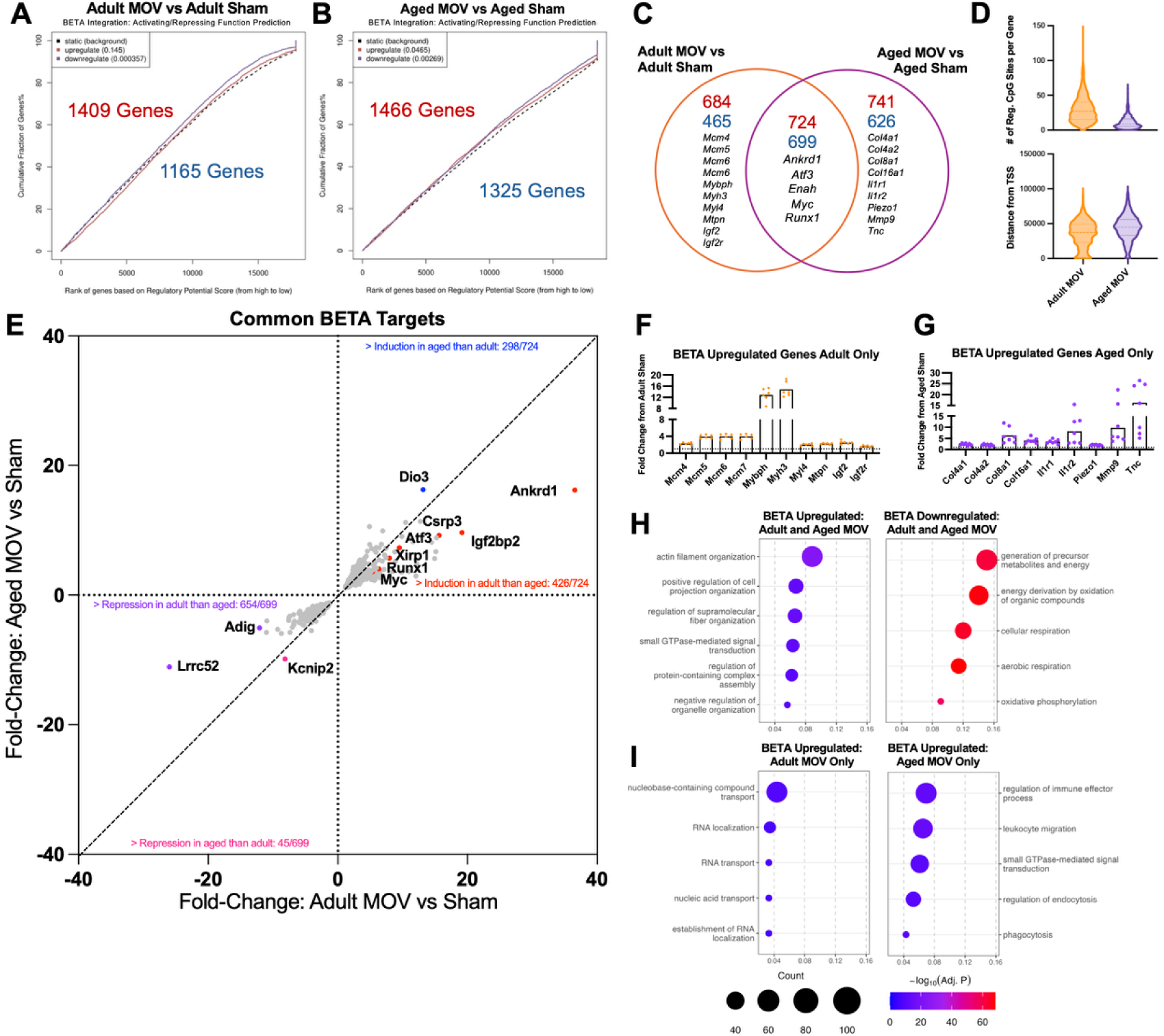
BETA integration analysis of myonuclear RRBS and bulk RNA-seq data comparing up- and downregulated genes relative to background (significance indicated by *p*-values in parentheses) for: (A) adult MOV vs adult sham, (B) aged MOV vs aged sham. (C) Venn diagram of genes regulated by methylation. (D) violin plots showing # of regulatory CpG sites for each BETA target and average absolute distance from the transcription start site. (E) X-Y plot showing fold changes of genes that are common BETA targets between adult MOV and aged MOV. (F) Plot of fold-changes of select BETA targets exclusive to adult MOV, (G) Plot of fold-changes of select BETA targets exclusive to aged MOV. (H) Top 5 enriched GO Biological Processes pathways from common upregulated and downregulated BETA targets. (I) Top 5 enriched GO Biological Processes pathways from upregulated BETA targets exclusive to adult MOV (left) and aged MOV (right). TSS = transcription start site. Reg = regulatory.

We next examined unique gene targets inferred to be regulated by DNA methylation during MOV in adult versus aged. There were 684 and 465 genes predicted to be up- or downregulated exclusively in adult MOV, respectively (Figure 7C). Adult MOV featured predicted methylation regulation of members of the minichromosome maintenance family: *Mcm4, Mcm5, Mcm6, Mcm7.* These genes are implicated in initiation of DNA replication in eukaryotes. Minichromosome gene regulation in adult myonuclei but not aged is provocative since it could be related to *de novo* resident myonuclear DNA synthesis observed during MOV-induced muscle growth^95–97^. Myosin binding protein H (*Mybph*) was upregulated and predicted to be controlled by methylation only in adult MOV (Figure 7F). *Mybph* is upregulated in peripheral artery disease (PAD) and the neurodegenerative disease amyotrophic lateral sclerosis (ALS)^98,99^, perhaps as a compensatory response. At the pathway level, BETA predicted genes exclusive to adult MOV corresponded to RNA and nucleic acid transport (Figure 7I, left). There were 741 up- and 626 downregulated genes predicted to be controlled by methylation exclusive to aged MOV (Figure 7C). These genes were related to ECM and inflammation/immune process pathways, with leukocyte migration and phagocytosis being the top enriched gene ontologies (Figure 7I, right). More specifically, aged muscle was highly enriched for the collagen genes *Col4a1, Col4a2, Col8a1,* and *Col16a1* and interleukin receptors *Il1r1* and *Il1r2* with MOV (Figure 7G). We previously reported robust induction of collagen and ECM remodeling genes in myonuclei during MOV in young mice (∼2 month old), but the age-specific methylome-transcriptome signature observed here may relate to impaired ECM adaptation during loading when aged^48^.

### Single myonucleus RNA-sequencing in adult versus aged muscle reveals age-dependent regulation of innervation and *Nr4a3* genes, which are linked to myonuclear DNA methylation

We next profiled the transcriptomes of 5,436 myonuclei (Figure 8A, 716-2,797 per experimental condition, Supplemental File 4). Initially, data from all experimental conditions were integrated and used to derive myonuclear clusters (Figure 8A). UMAP grouped by MOV versus sham, irrespective of age, are shown in Figure 8B. Four clusters were defined by myosin heavy chain gene expression (*Myh2, Myh1, Myh4*). These clusters were grouped together, termed “body myonuclei”, and comprised ∼90-95% of myonuclei in sham conditions (Figure 8C). Two of the seven myonuclear clusters were from specialized compartments such as the myotendinous junction (MTJ, ∼2.5%) and neuromuscular junction (NMJ, ∼0.6%). MTJ myonuclei were characterized by high expression of *Col22a1, Lama2, App,* and *Fras1,* and NMJ myonuclei by genes such as *Ache, Chrne, Ano4, Etv5*, *Vav3,* and *Col4a3* (Figure 8D). The remaining cluster was an “*Atf3*+” population which expanded by ∼15% in MOV groups compared to sham (Figure 8D, Supplemental Figure 4, Supplemental File 4).

**Figure 8.**
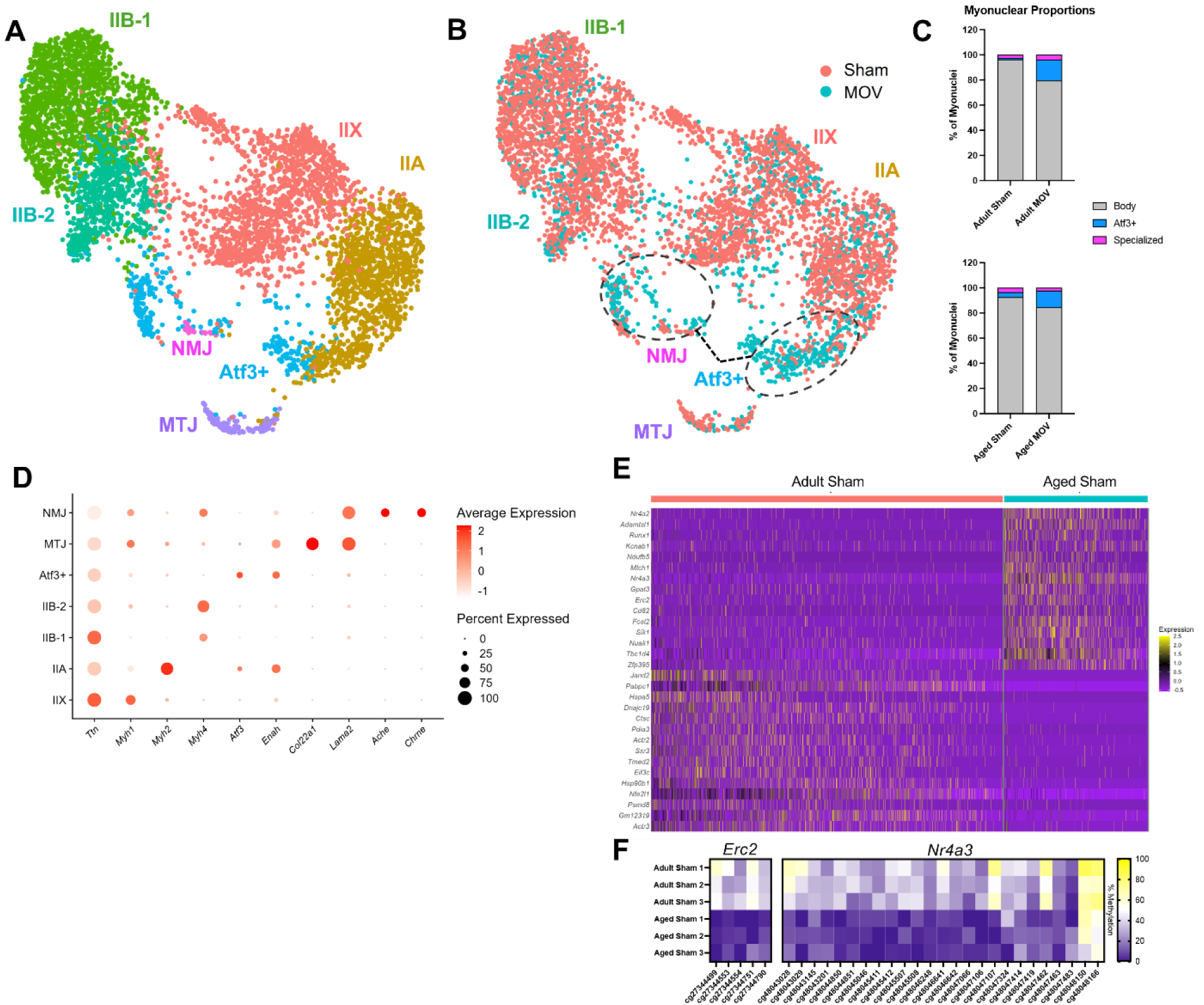
(A) Integrated UMAP of integrated all experimental conditions, labeled/colored by cluster. (B) Split UMAP of each experimental condition colored by cluster. (C) Percent composition of adult and aged myonuclei by cluster (D) Dot plot depicting top differentially marker genes for each myonuclear cluster. Dot size represents the percentage of nuclei expressing the gene. (E) Heatmap of top 20 up- and down-regulated genes between aged sham and adult sham. (F) Heatmap showing myonuclear DNA methylation percentage between aged sham and adult sham for select upregulated genes. NMJ = Neuromuscular Junction. MTJ = Myotendinous Junction. Body = Combined myosin expressing clusters.

We first defined the difference in aged sham versus adult sham myonuclei at single nucleus resolution to assess the effect of aging. We found enrichment of *Erc2, Nr4a3,* and *Runx1* in aged myonuclei, among other genes (Figure 8E). According to BETA, *Erc2* upregulation was associated exclusively with hypomethylated regulatory CpGs located in close proximity to the TSS in myonuclei (<360 bp, Figure 8F). Elevated *Erc2* was recently implicated in reduced muscle mass with aging and plays a role in organizing presynaptic active zones^100,101^. The *Nr4a* nuclear receptor family genes are among the most highly exercise responsive genes in human muscle, serving as a critical regulator of glucose and lipid metabolism^102–104^. Downregulation of *Nr4a* genes is deleterious for muscle mTORC1 signaling, ribosome biogenesis, and protein synthesis in muscle^102,103,105,106^. Upregulation of *Nr4a3* expression is typically associated with exercise adaptation and it is downregulated with inactivity in human muscle^102^. The link with aging is less well-known. Our bulk RNA-seq data do not show this family of genes to be altered with age; however, previous reports with snRNA-seq and snRNA-FISH show *Nr4a3* to be upregulated in aged tibialis anterior and gastrocnemius muscle^107^. Accordingly, *Nr4a3* was robustly hypomethylated in aged myonuclei relative to adult in our data (Figure 8F). Upregulation of *Nr4a3* with age could be specific to the ultra-fast glycolytic myosin types found in mice given *Nr4a3* is downregulated in aged human muscle, which is generally comprised of mixed slow and fast-oxidative myosin types^108^. Induction of *Runx1* is associated with muscle denervation during cachexia and aging in muscle^109–111^. *Runx1* expression alignes with observations of more denervated fibers with aging, specifically in fast-glycolytic fibers^111,112^. *Runx1* was not predicted to be regulated by BETA with aging alone, albeit there was considerable differential methylation (both hypo- and hypomethylation) at promoter and intron regions (Supplemental Figure 3).

### Age-dependent myonucleus-specific gene expression revealed by acute MOV in adult and aged muscle

Only one previous investigation has performed single myonucleus RNA-sequencing (smnRNA-seq) with MOV, and this was in young mice^113^. There are several notable differences in experimental design to consider between our study and theirs. First, in the previously published study, the myonuclear GFP labeling period was prior to and during MOV. This strategy captures newly fused satellite cells in addition to resident myonuclei. In the current study, resident myonuclei were labeled prior to MOV, thus excluding satellite cell-derived myonuclei during FANS-isolation. This difference in experimental design is important to consider given the prior study used 3-month-old mice, an age at which mice may still be undergoing developmental muscle growth and be reliant on satellite cell fusion and myonuclear accretion for initiating muscle hypertrophy compared to mice >4 months of age^18,114,115^. Our experiment was conducted in mature mice (>6 months). Second, in the prior study, the MOV period was 7 days versus our 3 days. We are evaluating the inherent ability of resident myonuclei to support hypertrophy largely independent from significant satellite cell fusion, which tends to occur later during MOV^33^. It is also worth mentioning that our smnRNA-seq was carried out on portions of the same muscles used for bulk RNA-seq and myonuclear RRBS, which enhances our ability to validate conclusions across assays.

With the modest number of myonuclei in our analysis, we first chose to broadly define how adult and aged myonuclei respond to MOV independent of cluster. Many of the top upregulated genes with MOV were common to adult and aged myonuclei: *Atf3, Ankrd1, Igf2bp2,* and *Creb5* (Figure 9A and 9B). Still, many genes were uniquely upregulated in adult MOV such as *Cd44, Pam,* and *Runx1*. *Cd44* is necessary for proper muscle regeneration, as *Cd44* knockout results in reduced muscle size and delayed recovery after muscle injury^116^. Perhaps higher *Cd44* in adult myonuclei with MOV contributes to age-specific muscle plasticity in a yet undefined way.

**Figure 9.**
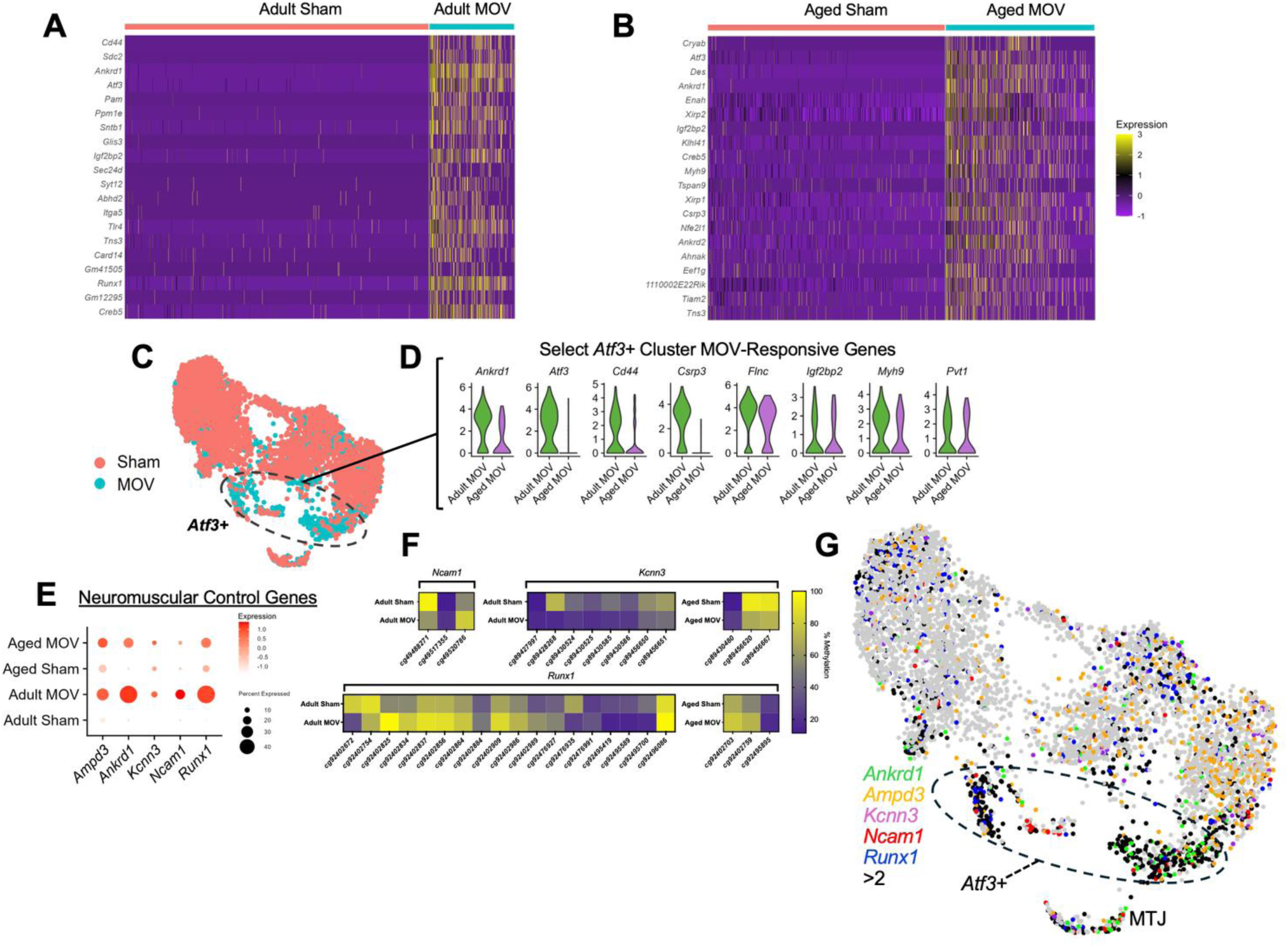
(A) Heatmap of top 20 upregulated genes between adult MOV and adult sham. (B) Heatmap of top 20 upregulated genes between aged MOV and aged sham. (C) UMAP grouped by MOV vs Sham. (D) Violin Plot showing expression of select genes upregulated in the Atf3+ population during MOV. In adult MOV, most genes are significant adj. *p* < 0.05. In aged MOV, no genes reached adj. *p* < 0.05 or were ns at *p* < 0.05. (E) Bubble plot showing expression of select genes involved in neuromuscular junction control. (F) Heatmap showing myonuclear DNA methylation percentage between MOV and sham conditions for neuromuscular control genes. (G) UMAP feature plot of neuromuscular control genes. Nuclei expressing >2 of the listed genes are colored black. ns = not significant.

We next focused on the *Atf3*+ population because it expands with MOV from ∼2% of myonuclei in sham to ∼15% of myonuclei in MOV in both ages (Figure 8C and 9C, Supplemental File 4). We performed cluster-level analysis on this population, and *Ankrd1, Atf3, Cd44, Crsp3, Flnc, Igf2bp2, Myh9,* and *Pvt1* were relatively more induced in adult versus aged MOV (Log_2_Fold changes; *Ankrd1*: 9.9 vs 4.9, *Atf3*: 10.1 vs 8.6; *Cd44*: 8.3 vs 4.0; *Csrp3*: 6.3 vs 2.05; *Flnc*: 3.75 vs 2.2; *Igf2bp2*: 5.7 vs 4; *Myh9*: 3.01 vs 3.14, *Pvt1:* 1.32 vs 0.88 in adult versus aged, respectively) (Figure 9D). Enrichment for several of these genes is similar to what is found in an *Atf3+* “sarcomere assembly” population defined previously during skeletal muscle development^107^ and in response to MOV in young mice^104^. Some of these same genes are enriched in resident myonuclei that migrate to sites of focal muscle fiber membrane damage^117^. Perhaps this emergent resident population with MOV are migrating “injury repair” myonuclei, and higher expression of these genes in adults contributes to superior sarcolemmal repair versus aged. The lncRNA *Pvt1,* which was upregulated during MOV in adult myonuclei but not in aged myonuclei, can interact with and stabilize *c-* and *n-Myc* (Figure 9D)^118^. These specific myonuclei may be part of an early myofibrillogenesis program that occurs during rapid developmental muscle growth (circa post-natal day 21) as well as MOV observed here, with a stronger response in adult versus aged. The induction of *Igf2bp2* (Insulin-like growth factor mRNA binding protein 2) in the *Atf3*+ population is intriguing. *Igf2bp2* binds *Igf2*, and *Igf2* was previously shown to be enriched in a myonuclear population derived from fused muscle satellite cells during MOV^113^. Although this *Igf2+* population was not detected in our dataset due to methodological differences, complementary expression of these two genes may reflect a collaboration between two myonuclear populations (resident and satellite-cell derived) to coordinate the response to MOV.

In *Atf3*-high myonuclei, we found upregulation of genes previously implicated in muscle innervation processes (*Ampd3, Ankrd1, Kcnn3, Ncam1, Runx1,* Figure 9E), and to a relatively greater extent in adult versus aged MOV. All of these genes were predicted to be regulated by methylation in myonuclei, with *Kcnn3, Ncam1, and Runx1* associated with myonuclear hypomethylation of promoter regions (Figure 9F). *Ampd3* and *Ankrd1* have methylation signatures at sites more distant from the TSS (shown in Supplemental File 3). When overlaid on the UMAP, these genes show high co-expression (black dots, >2 genes expressed in each nucleus) across the MOV responsive *Atf3*+ population, as well as MTJ myonuclei (Figure 9G)^109,110,119^. We did not identify myonuclei enriched for *H19*, *Igf2*, *Myh3* and *Myh8* (embryonic and neonatal myosin heavy chains) as observed previously during a longer duration of MOV in young mice^113^. The absence of these nuclei emphasizes that the emergence of these genes is a feature of recently fused satellite cells, which do not exist in our dataset (Supplemental File 4).

## Discussion

Aged muscle has a reduced capacity to grow in response to a hypertrophic stimulus^7–11^. Understanding the mechanisms underlying an age-associated loss of muscle plasticity is complicated by the syncytial nature of myofibers and the influence of non-muscle cell types that obscure myonuclear-specific changes. Parsing the roles of resident versus satellite cell-derived myonuclei is also important since resident myonuclei initiate the muscle hypertrophic process^14^. Taking advantage of the doxycycline-inducible genetically-modified HSA-GFP mouse model^12,15^, which enables the isolation of a high-purity population of resident myonuclei, in addition to myonuclear DNA methylation analysis and bulk RNA-sequencing, we evaluated the myonuclear molecular responses during the early phase of loading-induced muscle growth across several molecular layers. These datasets will serve as a resource for those at the intersection of skeletal muscle biology and aging research.

In sham mice, multi-omic integration revealed aged myonuclei experience broad changes in the transcriptome that is likely controlled by DNA methylation. Aging was associated with higher expression of ribosomal and mitochondrial gene expression and lower expression of ECM–related genes concomitant with changes to the DNA methylome in these same genes. Elevated ribosome gene regulation with age could be related to higher protein synthesis and dysregulated proteostasis that occurs during the muscle aging process^27^. A possible accumulation of dysfunctional ribosomes could also affect ribophagy, the autophagic degradation process for ribosomes, and compromise translational capacity in response to an anabolic stimulus^120^. Aged sham myonuclei have elevated expression of the transcription factor *Runx1* relative to young*. Runx1* is upregulated in denervated muscle^109,110^. Higher *Runx1* with age supports previous observations in aging humans suggesting an increased number of denervated fibers, specifically fast muscle fibers^119,121–123^. Altered innervation could contribute to a compromised ability to respond to a muscle growth stimulus. In previous work, an *Ampd3+* population of myonuclei appears in 30-month-old mice, which was posited to represent a dysfunctional denervated state given they were co-expressing pro-atrophy and proteasome-related genes^107^. Our data broadly align with a molecular signature of denervation from aging that is discernable at the epigenetic level in myonuclei.

By integrating myonuclear methylome and bulk transcriptome data, we provide the first detailed information on the myonuclear molecular landscape following a hypertrophic growth stimulus in adult versus aged mice. A similar number of genes displayed coordinated epigenome-transcriptome regulation after MOV in adult and aged muscle (∼1400 genes, 36% overlap). Our bulk RNA-sequencing show adult and aged muscle have a large conserved set of genes responsive to MOV at this early timepoint, including those involved in cytoskeletal organization and immune signaling along with suppression of oxidative phosphorylation pathways, as we have previously shown^36,48^. *miR-1* emerged as a key regulator of this metabolic reprogramming, being strongly repressed with MOV in both ages. With our inducible knockout or *miR-1* in muscle, we provide evidence that this myomiR is a powerful regulator of the skeletal muscle transcriptome and its repression explains some of the commonalities in the transcriptomes of adult and aged MOV, specifically as it relates to metabolism gene expression.

Although there were transcriptional profiles common to MOV-induced gene expression among adult and aged, aged muscle tended to have a blunted response. Our data suggest this could be attributed to a few factors: 1) the change in percent methylation of DM CpG sites in aged MOV were generally much lower than adult, 2) there are on average fewer regulatory CpG sites per target gene in aged, and 3) regulatory CpG sites are further from the transcription start site (or promoter region) in aged. Combined, these factors may lead to aged myonuclei having less control over each gene, potentially constraining the necessary “ramp up” of transcription to maximally adapt to an anabolic stimulus. To this point, aged muscle displayed a weaker induction of myonuclear enriched genes previously implicated in muscle mass regulation, including *Ankrd1, Mybph, Myc*, and *Runx1*. Instead, aged muscle showed a stronger induction of inflammatory, immune system, and cytokine-oriented genes with MOV. There are also numerous genes implicated in muscle mass regulation that were uniquely regulated in adult versus aged MOV muscle, such as minichromosome maintenance genes, that may contribute to differential adaptation between adult and aged.

Our smnRNA-seq data reinforce previous work showing expansion of an *Atf3*-enriched myonuclear subpopulation after overload^113^ - a pattern observed in both age groups. However, some genes enriched in this cluster generally exhibited weaker induction in aged than adult. This population has previously been thought to represent an early myofibrillogenesis or sarcomere assembly program that is similar to what occurs in post-natal muscle growth^113^. Adult muscle also featured stronger induction and epigenetic coordination of genes associated with neuromuscular remodeling: *Ampd3, Ankrd1, Kcnn3, Ncam1,* and *Runx1*. These observations, in context with our prior bulk myonuclear RNA-seq in young mice^48,107^ and alongside the observation that some of these genes are also upregulated by aging alone, raises a few possibilities: 1) acute MOV causes rapid neuromuscular junction (NMJ) remodeling that induces a de/re-innervation-like signature; 2) these genes are highly pleiotropic and act in a condition-dependent context to regulate muscle remodeling independent from NMJ disruption, or 3) all events are happening simultaneously. Nevertheless, the stronger induction of NMJ-related genes in adult versus aged MOV suggests an age-specific myonucleus-controlled neural contribution to the early hypertrophic process, which is further supported by our cellular deconvolution data.

There are a few limitations to our study that are worth considering. Our study only used male mice and we cannot confirm whether these results would be replicated in female mice. Additionally, we are capturing the early stage of the muscle response to MOV, where growth has not yet occurred, which may not necessarily reflect the molecular responses at later timepoints. Analyzing later time points and using different resistance-exercise models may be helpful for validating the molecular signatures we observe, as well as age-related differences in methylation/gene expression. Lastly, our smnRNA-seq is a limited dataset that did not allow us to fully evaluate the myonuclear transcriptome in high resolution; however, usage of the same muscles for all analyses increases the robustness of our conclusions, as does agreement with previously published smnRNA-seq and bulk myonuclear RNA-seq datasets from young animals^48,113^. Limitations aside, our integrated multimodal datasets provide unprecedented and detailed information on resident myonucleus-specific hypertrophic responses and could lead to new therapeutic targets for enhancing muscle adaptability in old age.

## Methods

### Animals

All animal procedures were approved by the University of Arkansas IACUC. Mice were housed in a temperature and humidity-controlled room, maintained on a 12:12-h light-dark cycle, and food and water were provided *ad libitum* throughout experimentation. At euthanasia (morning, ZT 1-5), animals were first deeply anesthetized with isoflurane and sacrificed *via* cervical dislocation. Adult (6-8 months of age) and old (24 months of age) male human skeletal actin reverse tetracycline transactivator - tetracycline response element histone 2B green fluorescent protein (HSA-rtTA^+/−^;TRE-H2B-GFP^+/−^, or HSA-GFP) mice were generated and genotyped as previously described by us^12,15^. HSA-GFP mice were treated with low-dose doxycycline (0.5 mg/ml doxycycline in drinking water with 2% sucrose) for 7 days to induce GFP labeling of myonuclei, followed by a washout period (normal drinking water) of at least 7 days. This strategy leads to the labeling of ∼95% of myonuclei with minimal off-target labeling of non-myonuclei. To knockout *miR-1* in adult mouse skeletal muscle, skeletal muscle-specific inducible *Mer-Cre-Mer* (HSA-MCM) mice were crossed with *miR-1-1^fl/fl^; miR-1-2^fl/fl^* (*miR-1^fl/fl^*) mice to produce HSA-MCM^+/-^; *miR-1-1^fl/fl^; miR-1-2^fl/fl^* mice (termed HSA-miR-1, KO)^40,124^. HSA-MCM^-/-^*; miR-1^fl/fl^* littermates mice served as controls (CON).

### Synergist Ablation Mechanical Overload Experiment

Synergist ablation mechanical overload (MOV) of the plantaris was performed as previously described by our lab group at a consistent daily interval (ZT 1-5)^12,48^. Briefly, synergist ablation is a surgical procedure that occurs while mice are under anesthesia. The surgery involves making an incision on the posterior aspect of the lower hindlimb, cutting of the Achilles tendon followed by careful removal of ∼30% the gastrocnemius–soleus complex while leaving the plantaris muscle and tendon intact. Sham surgery (control) involved all the steps of synergist ablation but no tendon is cut or muscle is removed. Following surgery mice resumed ambulatory cage activity. Euthanasia was performed 72 hours after MOV and both plantaris muscles were dissected and immediately flash frozen in liquid nitrogen. All muscle was used for downstream molecular analyses across four experimental conditions: adult sham (*n=*6), adult MOV (*n=*6), aged sham (*n=*6), aged MOV (*n=*7). The same cohort of animals were used to perform all experiments reported (Figure 1).

### Inducible Muscle-Specific miR-1 Knockout Experiment

Middle-aged (10-12 month-old) male HSA-miR-1 and *miR-1^fl/fl^* mice (*n*=3/group) were administered tamoxifen (2 mg/day) by intraperitoneal injection for five consecutive days, followed by an 8-week chase period. At euthanasia, KO and CON mice had all lower hindlimb muscle rapidly dissected, with one limb flash frozen for molecular analyses. Plantaris muscle was used for analyses.

### RNA Isolation, cDNA Synthesis, and Gene Expression Analysis

RNA was isolated from approximately one-third of each plantaris muscle (∼10mg) using TRIzol® Reagent (Sigma-Aldrich, St. Louis, MO, USA). Tissue was homogenized using zirconia beads and the Fisher Bead Mill (Fisher, Hampton, NH, USA). Following homogenization, RNA was isolated *via* phase separation by addition of chloroform and then centrifugation. The aqueous phase was transferred to a new sterile tube and further processed on spin columns according to manufacturer instructions using the Direct-zol Kit (Zymo Research, Irvine, CA, USA). RNA quality was checked on Agilent TapeStation with RNA screentape to confirm quality and purity. RNA integrity number (RIN) was >7 for all samples (8.3±0.7). For RT-qPCR of *miR-1* expression, cDNA was synthesized using the TaqMan MicroRNA Kit (4366596, Thermo Fisher Scientific, Waltham, MA). Gene expression of *miR-1* was analyzed by qPCR using TaqMan MicroRNA Assays (4427975, Thermo Fisher) as follows: *miR-1* (Assay ID 002222) and *U6* snRNA as the endogenous control for normalization (Assay ID 001973). The 2^-(ΔΔCt) method was used to calculate fold change.

### RNA Sequencing, Data Processing, and Statistical Analysis

RNA was sequenced by Novogene on an Illumina HiSeq using 150 bp paired-end sequencing, as we have previously done^48^. Raw FASTQ files were processed in Partek Flow. Alignment was performed using STAR 2.7.8a, quantified to annotation model mm39, filtered for features with a maximum of <5 counts, then normalization and statistical comparisons were performed with DESeq2. Genes with a false discovery rate (Benjamini–Hochberg method) adjusted *p*-value < 0.05 were identified as differentially expressed genes (DEGs). No fold-change cut offs were used. Pathway analyses were performed on up- and downregulated DEGs in R Studio (version 2025.09.0.387) with the 2025 gene ontology (GO) database as our cross reference (GO.db: Bioconductor version 3.21). We used all protein-coding genes detected in our RNA-sequencing dataset as our background correction for the pathway analysis^125^. For comparison of adult and old MOV RNA-seq data to myonuclear RNA-seq data during MOV, the list of upregulated genes from a previous study was used as a reference^48^. Figures were generated in GraphPad Prism version 10.6 for Mac OS X (GraphPad Software, La Jolla, CA) and RStudio.

### Digital Deconvolution of Cell Composition using Granulator

Cell type abundance was predicted from bulk RNA-sequencing data using Bioconductor R package Granulator (https://bioconductor.org/packages/release/bioc/html/granulator.html)^50^. By referencing single cell RNA-sequencing data, Granulator infers cell type abundance by modeling gene expression levels as weighted sums of the cell-type specific expression profiles. We used skeletal muscle single-cell RNA-seq data from 4-day murine plantaris tenotomy data from Zhang et al^50^. The publicly available datasets (10X Genomics .h5 files) were downloaded from GEO (GSE232257), reanalyzed with Seurat, and cell clusters were identified using the exact parameters outlined in the initial publication. Normalized gene expression matrices of each cell type served as a reference gene expression matrix was integrated with normalized counts from our bulk RNA-sequencing data, and cell proportions were predicted by Granulator. Statistical analyses of predicted cell-type proportions were analyzed by 2-way ANOVA to determine main effects of age, MOV, or interactions. Significance levels were established *a priori* at *p* < 0.05.

### Fluorescent Activated Nuclear Sorting (FANS)

Myonuclei were isolated *via* Fluorescent Activated Nuclear Sorting (FANS) on a MACSQuant Tyto Cell Sorter (Miltenyi Biotec, Bergisch Gladbach, Germany). For myonuclear DNA methylation experiments, approximately one-half of each plantaris was used (∼15mg). Muscle was placed in a small glass beaker a sucrose-based buffer mimicking physiological conditions (5 mM PIPES, 85mM KCl, 1mM CaCl_2_, 5% sucrose, 2X HALT Protease inhibitors, and 0.25% NP-40), pulsed with 4 μL of propidium iodide, then minced with scissors until a slurry. The nuclear suspension was then transferred to glass Dounce for further manual homogenization, then strained through a 20-μm MACSQuant pre-separation filter directly into MACSQuant Tyto regular-speed sorting cartridge (Cat #: 130-104-791). For single myonucleus RNA-sequencing experiments, remaining muscle (∼2-3mg) of plantaris from three mice per condition were pooled in homogenization buffer (500 µL HEPES [1 M], 3 mL KCl [1 M], 250 µL spermidine [100 mM], 750 µL spermine tetrahydrochloride [10 mM], 10 mL EDTA [10 mM], 250 µL EGTA [100 mM], 2.5 mL MgCl [100 mM], 5.13 g sucrose) with 0.2U/µL RNAse inhibitors (Protector RNase Inhibitor, Millipore Sigma, Burlington, MA, USA). The muscle was minced in buffer with scissors on ice in a low-bind 1.5 mL tube, dounced ∼20 times with a plastic pestle with a gentle twist at the bottom, then strained through a 20-μm MACSQuant pre-separation filter (Cat #: 130-101-812) directly into MACSQuant Tyto high-speed sorting cartridge (Cat #: 130-121-549). In both experiments, FANS gating was established to exclude debris and identify nuclei positive for both GFP (intrinsic myonuclear label) and PI, then sorted directly into the respective buffer for downstream analyses. Myonuclei for DNA methylation analyses were sorted into ATL buffer and proteinase K (Qiagen) for genomic DNA purification. Myonuclei for smnRNA-sequencing analyses were sorted into PBS/1% BSA/RNase inhibitors.

### Myonuclear Genomic DNA Isolation, Reduced Representation Bisulfite Sequencing, and Analysis

DNA isolation was carried out according to the previously described by us with minor adjustments^10^. Briefly, using the QIAamp DNA micro kit (Qiagen) myonuclei sorted into buffer ATL and proteinase K were incubated for a minimum of 4 hours at 56°C. DNA binding to the column was conducted using 1 µg of carrier RNA, and washes and centrifugations were carried out according to the manufacturer’s instructions. DNA was eluted in 12 µl of nuclease-free H_2_O, quality checked on the Agilent Tapestation with the genomic DNA (gDNA) screen tape and placed in -20°C until later analyses. Low-input Msp1 Reduced Representation Bisulfite Sequencing (RRBS) was performed by Zymo Research using 5 ng of gDNA that was generally >40,000 base pairs (bp) in length. Some samples did not reach the minimum gDNA mass requirement for RRBS and could not be included in downstream analysis. Quality control and adapter sequence trimming were performed using FastQC and Cutadapt, respectively as parts of the Trim Galore wrapper. Low-quality base calls (Phred score <20) were removed prior to trimming adapter sequences. Bismark aligner was used to align the sequence reads to the bisulfite-converted mm39 genome prior to data processing. Coverage (.cov) ds produced from Bismark aligner were used for data analysis in the methylKit R package, with a minimum reads cut off of >10x coverage per CpG site across all samples and a minimum base coverage of 1 per sample, as previously described^2,72,126^. Percent methylation and percent differential methylation were then obtained from methylKit following analysis. Differentially methylated sites were defined as q-value of <0.05 and >10% methylation difference. During the initial stage of analysis, we found a few samples with extreme deviation from experimental conditions and potentially skewing statistical analyses. We confirmed this was not due to technical errors in tissue process or labeling. We ran a series of diagnostics on the annotated promoter matrices to assess whether any samples should be considered for exclusion. This included: 1) PCA with group centroid overlays, to visualize variance and clustering structure, 2) Centroid distance calculations, to quantify how far each sample is from the center of its group, 3) Multivariate dispersion testing, to assess within-group spread in an unsupervised way. Based on these results, we identified five samples that clearly fell outside the expected distribution across multiple comparisons, showing extreme distances from their respective group centroids, visually separated from their cohorts, and in some cases clustered with the other experimental condition (e.g. MOV vs Sham). Given the magnitude of variance in these samples and the uncertainty of how this variance arose, we excluded these samples from analysis. This left us with samples sizes for each condition of: adult sham (n=3), adult MOV (n=4), aged sham (n=3), aged MOV (n=3).

### BETA integration Pathway Analysis

Integration of differential DNA methylation and gene expression data was performed using BETA basic (Binding and Expression Target Analysis, v1.0.7). This software that provides an integrated analysis of transcription-factor binding to genomic DNA and transcript abundance using chromatin immunoprecipitation sequencing (ChIP-seq) and transcriptomics (RNA-seq) datasets^84^. BETA models the effect of regulatory elements using a natural-log function of their distance to the transcription start site (TSS) to calculate a regulatory potential score^127^. This method for integration of RRBS and RNA-seq data using BETA is consistent with our previous publications^36,72^. In this study, differentially methylated CpG sites derived from RRBS were formatted as BED “peak” files and used as the regulatory-element input, while differentially expressed genes (adjusted p < 0.05) from bulk RNA-seq were used as the expression input. Gene annotations were derived from the mm39 UCSC GTF converted to BED format. The BETA basic was executed with the following command: “-k BSF -- gname2 --df 0.05 --pn 100000 -c 0.05”. These parameters specify the binding-site kernel function, use of gene symbols, a 5% FDR cutoff for both methylation and expression data, and 100,000 permutations for robust significance estimation. CpG-associated peaks within 100 kb of a gene’s TSS were included in the calculation of the regulatory potential through the following equation:

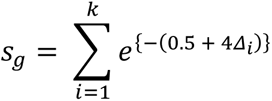

In this formulation, *k* represents the number of CpG-associated peaks (CpG islands) linked to gene *g*, and *Δ_i_* represents the distance of each element from the TSS, inversely scaled so that elements nearer the TSS exert greater influence on the regulatory potential score (*s_g_*). Genes were classified as activated, repressed, or non-targeted according to the direction of their expression change relative to the predicted regulatory potential. It is also worth mentioning not all CpG-associated peaks predicted to regulate gene activation or repression are found to be differentially methylated, and vice versa.

### Single Myonucleus RNA Sequencing Library Preparation and Analysis

For this analysis, plantaris muscle from 3 samples were pooled per condition for analysis – MOV versus sham in adult and aged (*n=*4 samples). GFP/PI+ myonuclei were isolated via FANS on our MACSQuant Tyto Sorter and sorted directly into 36 µl of PBS/1% BSA/RNase inhibitors to minimize dilution. This was the final analysis performed from the same cohort of mice, so at this stage we were limited to ∼2-3 mg of plantaris muscle per mouse. Combined with more gentle tissue dissociation methods compared to isolation of myonuclei for genomic DNA isolation (plastic pestle vs glass Dounce) in effort to preserve nuclei integrity, the resulting nuclei yield per unit mass of tissue mass was lower than the maximum input into the 10X chromium device. We combined 36.6 µl of the myonuclei suspension with 28.4 µl of reverse transcription master mix (MM). The 65 µl myonuclei + MM solution was loaded into the GEM-X 3’ chip for GEM formation, then the library was prepared using the Single Cell 3’ Reagent Kit v4 according to the manufacturer’s protocol. Following library construction, libraries were sequenced on the Illumina Nova NextSeq X Plus System by Novogene to 200 million reads per sample. Raw FASTQ files were imported into Cell Ranger 9.0 for alignment to reference transcriptome, and report of UMIs/reads/nuclei barcodes. h5 files were exported and imported into Python environment, where ambient RNA correction tool CellBender was used^128,129^. Processed files were imported into Seurat v5^130^, and data quality control was performed by removing nuclei with <200 UMIs, nuclei with >5% of mitochondrial reads, and genes expressed in <3 nuclei. Datasets were normalized using the SCTransform() command. Dimensionality reduction was performed using the RunPCA(), RunUMAP(), FindNeighbors(), FindClusters() commands. Doublet removal was performed using DoubletFinder(). All objects were then integrated using SelectIntegrationFeatures() from the top 3000 variable features and PreSCTIntegration(). Integration anchors were generated using the FindIntegrationAnchors() command, inputting the identified integration features to the “anchor.features” parameter, and specifying the “normalization.method” parameter as “SCT”. Datasets were then integrated by supplying the anchors to the IntegrateData() command, specifying the “normalization.method” parameter as “SCT”. A final dimensionality reduction was performed using RunPCA(), FindNeighbors(), FindClusters(), RunUMAP() with number of PCs set to 30 and resolution of 0.5. Data were then normalized and scaled with JoinLayers(), NormalizedData(), and ScaleData(). Clusters were visualized with DimPlot(). Gene expression visualization of clusters was performed using DoHeatmap(). Feature plots were generated using the FeaturePlot() function. DotPlots were created using the DotPlot() function. Violin plots were created using the VlnPlot() command. Input features for Heatmaps and Dotplots were either manually selected, or generated through the FindAllMarkers() or FindMarkers() functions. Differentially expressed genes between clusters were identified using min.pct = 0.25, Log_2_FC > 0.25, with adjusted *p*-value (FDR) < 0.05. Differentially expressed genes between specific experimental conditions were identified as min.pct = 0.1, Log_2_FC > 0.5, with adjusted *p*-value (FDR) < 0.05.

## Author Contributions

P.J.K. and K.A.M. conceived the study. P.J.K., R.G.J., A.R.C., F.M., and A.I. performed experiments and/or analysis. K.A.M, N.P.G., and J.J.M provided resources. P.J.K and K.A.M wrote the manuscript draft with input from A.I., J.J.M., N.P.G., and Y.W. All authors reviewed and approved the manuscript.

## Acknowledgements

Thank you to C. Brooks Mobley, PhD, for assistance with mouse breeding and colony management. The graphical abstract and experimental design figures were generated using BioRender.

## Data Availability

Bulk RNA-seq, RRBS, and smnRNA-seq data will be deposited in the Gene Expression Omnibus (GEO) database. Previously published bulk myonuclear RNA-seq MOV data are available in GSE213406^48^.

## Ethics Statement

All animal procedures were approved by the Institutional Animal Care and Use Committee of the University of Arkansas.

## Consent

The authors have nothing to report.

## Conflicts of Interest

Y.W. is the founder of MyoAnalytics LLC. The remaining authors have no other competing interests to declare.

**Supplemental Figure 1.** (A) Venn diagram comparing overlap in upregulated differentially expressed genes between adult MOV, aged MOV, and miR-1 KO. B) Venn diagram comparing overlap in downregulated differentially expressed genes between adult MOV, aged MOV, and miR-1 KO. C) AGO2 eCLIP-seq miR-1 binding peaks for genes ADP-ribosylation factor 4 (*Arf4*), antizyme inhibitor 1 (*Azin1*), and protein tyrosine phosphatase non-receptor type 1 (*Ptpn1*), from Ismaeel et al. *Molecular Metabolism*, 2025. Complete list of overlapping genes are available in Supplementary File 1.

**Supplemental Figure 2.** (A) BETA integration analysis of myonuclear RRBS and bulk RNA-sequencing data from aged sham vs adult sham comparing up- and downregulated genes relative to background (significance indicated by *p*-values in parentheses). (B&C) Plots showing top 10 up- and downregulated GO process pathways (biological processes, cellular component, molecular function) from genes identified to have regulation predicted by BETA integration.

**Supplemental Figure 3.** (A) Plot showing distance of BETA predicted regulatory CpG sites from TSS and regulatory score for *Myc*. (B) Heatmap of all differentially methylated CpG sites for *Myc* during MOV. (C) Table of additional muscle growth-related genes and number of BETA regulatory CpG sites and average distance from transcription start site (TSS).

**Supplemental Figure 4.** (A) Feature plots of select canonical muscle identity and fiber type markers: *Ttn, Myh1, Myh2, Myh4.* MTJ-specific markers: *Col22a1, Lama2, App.* NMJ-specific markers: *Etv5, Vav3, Ache*.

## References

1. Cartee GD, Hepple RT, Bamman MM, Zierath JR. Exercise Promotes Healthy Aging of Skeletal Muscle. Cell Metabolism. 2016;23(6):1034–1047. doi:10.1016/j.cmet.2016.05.007

2. Roberts MD, McCarthy JJ, Hornberger TA, et al. Mechanisms of mechanical overload-induced skeletal muscle hypertrophy: current understanding and future directions. Physiological Reviews. 2023;103(4):2679–2757. doi:10.1152/physrev.00039.2022

3. Chambers TL, Murach KA. A history of omics discoveries reveals the correlates and mechanisms of loading-induced hypertrophy in adult skeletal muscle. 2024 CaMPS young investigator award invited review. American Journal of Physiology-Cell Physiology. 2025;328(5):C1535–C1557. doi:10.1152/ajpcell.00968.2024

4. Hurst C, Robinson SM, Witham MD, et al. Resistance exercise as a treatment for sarcopenia: prescription and delivery. Age Ageing. 2022;51(2):afac003. doi:10.1093/ageing/afac003

5. McKendry J, Currier BS, Lim C, Mcleod JC, Thomas ACQ, Phillips SM. Nutritional Supplements to Support Resistance Exercise in Countering the Sarcopenia of Aging. Nutrients. 2020;12(7):2057. doi:10.3390/nu12072057

6. Law TD, Clark LA, Clark BC. Resistance Exercise to Prevent and Manage Sarcopenia and Dynapenia. Annuaul Review of Gerontology and Geriatrics. 2016;36(1):205–228. doi:10.1891/0198-8794.36.205

7. Greig CA, Gray C, Rankin D, et al. Blunting of adaptive responses to resistance exercise training in women over 75y. Experimental Gerontology. 2011;46(11):884–890. doi:10.1016/j.exger.2011.07.010

8. Welle S, Totterman S, Thornton C. Effect of age on muscle hypertrophy induced by resistance training. J Gerontol A Biol Sci Med Sci. 1996;51(6):M270–275. doi:10.1093/gerona/51a.6.m270

9. Kirby TJ, Lee JD, England JH, Chaillou T, Esser KA, McCarthy JJ. Blunted hypertrophic response in aged skeletal muscle is associated with decreased ribosome biogenesis. J Appl Physiol. 2015;119(4):321–327. doi:10.1152/japplphysiol.00296.2015

10. Dungan CM, Brightwell CR, Wen Y, et al. Muscle-Specific Cellular and Molecular Adaptations to Late-Life Voluntary Concurrent Exercise. Function. 2022;3(4):zqac027. doi:10.1093/function/zqac027

11. Nolt GL, Keeble AR, Wen Y, et al. Inhibition of p53-MDM2 binding reduces senescent cell abundance and improves the adaptive responses of skeletal muscle from aged mice. GeroScience. 2024;46(2):2153–2176. doi:10.1007/s11357-023-00976-2

12. Von Walden F, Rea M, Mobley CB, et al. The myonuclear DNA methylome in response to an acute hypertrophic stimulus. Epigenetics. 2020;15(11):1151–1162. doi:10.1080/15592294.2020.1755581

13. Pacheco C, Felipe SM da S, Soares MMD de C, et al. A compendium of physical exercise-related human genes: an ’omic scale analysis. Biol Sport. 2018;35(1):3–11. doi:10.5114/biolsport.2018.70746

14. Koopmans PJ, Zwetsloot KA, Murach KA. Going nuclear: Molecular adaptations to exercise mediated by myonuclei. Sports Medicine and Health Science. 2023;5(1):2–9. doi:10.1016/j.smhs.2022.11.005

15. Iwata M, Englund DA, Wen Y, et al. A novel tetracycline-responsive transgenic mouse strain for skeletal muscle-specific gene expression. Skelet Muscle. 2018;8(1):33. doi:10.1186/s13395-018-0181-y

16. Burke BI, Ismaeel A, Walden F von, Murach KA, McCarthy JJ. Response to: Counterpoint to: The Utility of the Rodent Synergist Ablation Model in Identifying Molecular and Cellular Mechanisms of Skeletal Muscle Hypertrophy. American Journal of Physiology-Cell Physiology. doi:10.1152/ajpcell.00418.2024

17. Lee JD, Fry CS, Mula J, et al. Aged Muscle Demonstrates Fiber-Type Adaptations in Response to Mechanical Overload, in the Absence of Myofiber Hypertrophy, Independent of Satellite Cell Abundance. J Gerontol A Biol Sci Med Sci. 2016;71(4):461–467. doi:10.1093/gerona/glv033

18. McCarthy JJ, Mula J, Miyazaki M, et al. Effective fiber hypertrophy in satellite cell-depleted skeletal muscle. Development. 2011;138(17):3657–3666. doi:10.1242/dev.068858

19. Clemens Z, Sivakumar S, Pius A, et al. The biphasic and age-dependent impact of klotho on hallmarks of aging and skeletal muscle function. Suh Y, Tyler JK, Remondini D, eds. eLife. 2021;10:e61138. doi:10.7554/eLife.61138

20. Ashburner M, Ball CA, Blake JA, et al. Gene Ontology: tool for the unification of biology. Nat Genet. 2000;25(1):25–29. doi:10.1038/75556

21. Ge SX, Son EW, Yao R. iDEP: an integrated web application for differential expression and pathway analysis of RNA-Seq data. BMC Bioinformatics. 2018;19(1):534. doi:10.1186/s12859-018-2486-6

22. Stec MJ, Mayhew DL, Bamman MM. The effects of age and resistance loading on skeletal muscle ribosome biogenesis. J Appl Physiol (1985). 2015;119(8):851-857. doi:10.1152/japplphysiol.00489.2015

23. Joseph GA, Wang, SX, Jacobs, CE, et al. Partial Inhibition of mTORC1 in Aged Rats Counteracts the Decline in Muscle Mass and Reverses Molecular Signaling Associated with Sarcopenia. Molecular and Cellular Biology. 2019;39(19):e00141–19. doi:10.1128/MCB.00141-19

24. Kaiser MS, Milan G, Ham DJ, et al. Dual roles of mTORC1-dependent activation of the ubiquitin-proteasome system in muscle proteostasis. Commun Biol. 2022;5(1):1–18. doi:10.1038/s42003-022-04097-y

25. Shavlakadze T, Zhu J, Wang S, et al. Short-term Low-Dose mTORC1 Inhibition in Aged Rats Counter-Regulates Age-Related Gene Changes and Blocks Age-Related Kidney Pathology. The Journals of Gerontology: Series A. 2018;73(7):845–852. doi:10.1093/gerona/glx249

26. Zarzycka W, Kobak KA, King CJ, Peelor FF, Miller BF, Chiao YA. Hyperactive mTORC1/4EBP1 signaling dysregulates proteostasis and accelerates cardiac aging. GeroScience. 2025;47(2):1823–1836. doi:10.1007/s11357-024-01368-w

27. O’Reilly CL, Bodine SC, Miller BF. Current limitations and future opportunities of tracer studies of muscle ageing. The Journal of Physiology. 2025;603(1):7–15. doi:10.1113/JP285616

28. Jang YC, Van Remmen H. Age-associated alterations of the neuromuscular junction. Experimental Gerontology. 2011;46(2):193–198. doi:10.1016/j.exger.2010.08.029

29. Abbott CB, Lawrence MM, Kobak KA, et al. A Novel Stable Isotope Approach Demonstrates Surprising Degree of Age-Related Decline in Skeletal Muscle Collagen Proteostasis. Function. 2021;2(4):zqab028. doi:10.1093/function/zqab028

30. Kragstrup TW, Kjaer M, Mackey AL. Structural, biochemical, cellular, and functional changes in skeletal muscle extracellular matrix with aging. Scandinavian Journal of Medicine & Science in Sports. 2011;21(6):749–757. doi:10.1111/j.1600-0838.2011.01377.x

31. Lawrence MM, Abbott C, Peelor III FF, Lopes EBP, Griffin TM, Miller BF. Determining Resistance to Protein Turnover in Aged Skeletal Muscle Collagen Using a Novel Stable Isotope Timecourse Approach. The FASEB Journal. 2020;34(S1):1–1. doi:10.1096/fasebj.2020.34.s1.01802

32. Garg K, Boppart MD. Influence of exercise and aging on extracellular matrix composition in the skeletal muscle stem cell niche. Journal of Applied Physiology. 2016;121(5):1053–1058. doi:10.1152/japplphysiol.00594.2016

33. Murach KA, Peck BD, Policastro RA, et al. Early satellite cell communication creates a permissive environment for long-term muscle growth. iScience. 2021;24(4):102372. doi:10.1016/j.isci.2021.102372

34. Baumert P, Mäntyselkä S, Schönfelder M, et al. Skeletal muscle hypertrophy rewires glucose metabolism: An experimental investigation and systematic review. J Cachexia Sarcopenia Muscle. 2024;15(3):989–1002. doi:10.1002/jcsm.13468

35. Wackerhage H, Vechetti IJ, Baumert P, et al. Does a Hypertrophying Muscle Fibre Reprogramme its Metabolism Similar to a Cancer Cell? Sports Med. 2022;52(11):2569–2578. doi:10.1007/s40279-022-01676-1

36. Ismaeel A, Thomas NT, McCashland M, et al. Coordinated Regulation of Myonuclear DNA Methylation, mRNA, and miRNA Levels Associates With the Metabolic Response to Rapid Synergist Ablation-Induced Skeletal Muscle Hypertrophy in Female Mice. Function (Oxf). 2023;5(1):zqad062. doi:10.1093/function/zqad062

37. Valentino T, Figueiredo VC, Mobley CB, McCarthy JJ, Vechetti Jr IJ. Evidence of myomiR regulation of the pentose phosphate pathway during mechanical load-induced hypertrophy. Physiological Reports. 2021;9(23):e15137. doi:10.14814/phy2.15137

38. Murach KA, Mobley CB, Zdunek CJ, et al. Muscle memory: myonuclear accretion, maintenance, morphology, and miRNA levels with training and detraining in adult mice. J Cachexia Sarcopenia Muscle. 2020;11(6):1705–1722. doi:10.1002/jcsm.12617

39. Karagkouni D, Paraskevopoulou MD, Chatzopoulos S, et al. DIANA-TarBase v8: a decade-long collection of experimentally supported miRNA–gene interactions. Nucleic Acids Res. 2018;46(Database issue):D239–D245. doi:10.1093/nar/gkx1141

40. Ismaeel A, Peck BD, Montgomery MM, et al. microRNA-1 Regulates Metabolic Flexibility by Programming Adult Skeletal Muscle Pyruvate Metabolism. Molecular Metabolism. Published online June 7, 2025:102182. doi:10.1016/j.molmet.2025.102182

41. Vechetti IJ, Peck BD, Wen Y, et al. Mechanical overload-induced muscle-derived extracellular vesicles promote adipose tissue lipolysis. FASEB J. 2021;35(6):e21644. doi:10.1096/fj.202100242R

42. Fei S, Rule BD, Godwin JS, et al. miRNA-1 regulation is necessary for mechanical overload-induced muscle hypertrophy in male mice. Physiological Reports. 2025;13(1):e70166. doi:10.14814/phy2.70166

43. Nakai W, Kondo Y, Saitoh A, Naito T, Nakayama K, Shin HW. ARF1 and ARF4 regulate recycling endosomal morphology and retrograde transport from endosomes to the Golgi apparatus. MBoC. 2013;24(16):2570–2581. doi:10.1091/mbc.e13-04-0197

44. Adarska P, Wong-Dilworth L, Bottanelli F. ARF GTPases and Their Ubiquitous Role in Intracellular Trafficking Beyond the Golgi. Front Cell Dev Biol. 2021;9. doi:10.3389/fcell.2021.679046

45. Sesen J, Martinez T, Busatto S, et al. AZIN1 level is increased in medulloblastoma and correlates with c-Myc activity and tumor phenotype. Journal of Experimental & Clinical Cancer Research. 2025;44(1):56. doi:10.1186/s13046-025-03274-1

46. Yu J, Zhang C, Zhang Q, et al. AZIN1-dependent polyamine synthesis accelerates tumor cell cycle progression and impairs effector T-cell function in osteosarcoma. Cell Death Dis. 2025;16(1):310. doi:10.1038/s41419-025-07640-x

47. Xie L, Qi H, Tian W, Bu S, Wu Z, Wang H. High-expressed PTPN1 promotes tumor proliferation signature in human hepatocellular carcinoma. Heliyon. 2023;9(9). doi:10.1016/j.heliyon.2023.e19895

48. Murach KA, Liu Z, Jude B, et al. Multi-transcriptome analysis following an acute skeletal muscle growth stimulus yields tools for discerning global and MYC regulatory networks. Journal of Biological Chemistry. 2022;298(11). doi:10.1016/j.jbc.2022.102515

49. Newman AM, Steen CB, Liu CL, et al. Determining cell type abundance and expression from bulk tissues with digital cytometry. Nat Biotechnol. 2019;37(7):773–782. doi:10.1038/s41587-019-0114-2

50. Zhang L, Saito H, Higashimoto T, et al. Regulation of muscle hypertrophy through granulin: Relayed communication among mesenchymal progenitors, macrophages, and satellite cells. Cell Reports. 2024;43(4):114052. doi:10.1016/j.celrep.2024.114052

51. Chen W, Datzkiw D, Rudnicki MA. Satellite cells in ageing: use it or lose it. Open Biol. 2020;10(5):200048. doi:10.1098/rsob.200048

52. Snijders T, Verdijk LB, Smeets JSJ, et al. The skeletal muscle satellite cell response to a single bout of resistance-type exercise is delayed with aging in men. Age (Dordr). 2014;36(4):9699. doi:10.1007/s11357-014-9699-z

53. Walker DK, Fry CS, Drummond MJ, et al. Pax7+ Satellite Cells in Young and Older Adults following Resistance Exercise. Muscle Nerve. 2012;46(1):51–59. doi:10.1002/mus.23266

54. Proietti D, Giordani L, De Bardi M, et al. Activation of skeletal muscle-resident glial cells upon nerve injury. JCI Insight. 2021;6(7):e143469, 143469. doi:10.1172/jci.insight.143469

55. Thomas NT, Brightwell CR, Owen AM, et al. Satellite cells choreograph an immune cell-fibrogenic cell circuit during mechanical loading in geriatric skeletal muscle. PNAS Nexus. 2025;4(9):pgaf236. doi:10.1093/pnasnexus/pgaf236

56. Murach KA, Vechetti IJ Jr, Van Pelt DW, et al. Fusion-Independent Satellite Cell Communication to Muscle Fibers During Load-Induced Hypertrophy. Function. 2020;1(1):zqaa009. doi:10.1093/function/zqaa009

57. Fry CS, Kirby TJ, Kosmac K, McCarthy JJ, Peterson CA. Myogenic Progenitor Cells Control Extracellular Matrix Production by Fibroblasts during Skeletal Muscle Hypertrophy. Cell Stem Cell. 2017;20(1):56–69. doi:10.1016/j.stem.2016.09.010

58. Moorwood C, Barton ER. Caspase-12 ablation preserves muscle function in the mdx mouse. Hum Mol Genet. 2014;23(20):5325–5341. doi:10.1093/hmg/ddu249

59. Li S, Karri D, Sanchez-Ortiz E, et al. Sema3a-Nrp1 Signaling Mediates Fast-Twitch Myofiber Specificity of Tw2+ Cells. Developmental Cell. 2019;51(1):89–98.e4. doi:10.1016/j.devcel.2019.08.002

60. Liu N, Garry GA, Li S, et al. A Twist2-dependent progenitor cell contributes to adult skeletal muscle. Nat Cell Biol. 2017;19(3):202–213. doi:10.1038/ncb3477

61. Gaulton N, Wakelin G, Young LV, et al. Twist2-expressing cells reside in human skeletal muscle and are responsive to aging and resistance exercise training. The FASEB Journal. 2022;36(12):e22642. doi:10.1096/fj.202201349RR

62. Cameron A, Wakelin G, Gaulton N, et al. Identification of underexplored mesenchymal and vascular-related cell populations in human skeletal muscle. American Journal of Physiology-Cell Physiology. 2022;323(6):C1586–C1600. doi:10.1152/ajpcell.00364.2022

63. Peck BD, Murach KA, Walton RG, et al. A muscle cell-macrophage axis involving matrix metalloproteinase 14 facilitates extracellular matrix remodeling with mechanical loading. The FASEB Journal. 2022;36(2):e22155. doi:10.1096/fj.202100182RR

64. Schoeps B, Frädrich J, Krüger A. Cut loose TIMP-1: an emerging cytokine in inflammation. Trends in Cell Biology. 2023;33(5):413–426. doi:10.1016/j.tcb.2022.08.005

65. Qin CC, Liu YN, Hu Y, Yang Y, Chen Z. Macrophage inflammatory protein-2 as mediator of inflammation in acute liver injury. World J Gastroenterol. 2017;23(17):3043–3052. doi:10.3748/wjg.v23.i17.3043

66. De Plaen IG, Han XB, Liu X, Hsueh W, Ghosh S, May MJ. Lipopolysaccharide induces CXCL2/macrophage inflammatory protein-2 gene expression in enterocytes via NF-κB activation: independence from endogenous TNF-α and platelet-activating factor. Immunology. 2006;118(2):153–163. doi:10.1111/j.1365-2567.2006.02344.x

67. Edman S, Jones III RG, Jannig PR, et al. The 24-hour molecular landscape after exercise in humans reveals MYC is sufficient for muscle growth. EMBO reports. Published online October 31, 2024:1–28. doi:10.1038/s44319-024-00299-z

68. Jones RGI, von Walden F, Murach KA. Exercise-Induced MYC as an Epigenetic Reprogramming Factor That Combats Skeletal Muscle Aging. Exercise and Sport Sciences Reviews. 2024;52(2):63. doi:10.1249/JES.0000000000000333

69. Alway SE. Overload-Induced C-Myc Oncoprotein Is Reduced in Aged Skeletal Muscle. J Gerontol A Biol Sci Med Sci. 1997;52A(4):B203-B211. doi:10.1093/gerona/52A.4.B203

70. Rivas DA, Lessard SJ, Rice NP, et al. Diminished skeletal muscle microRNA expression with aging is associated with attenuated muscle plasticity and inhibition of IGF-1 signaling. FASEB J. 2014;28(9):4133–4147. doi:10.1096/fj.14-254490

71. Brook MS, Wilkinson DJ, Mitchell WK, et al. Synchronous deficits in cumulative muscle protein synthesis and ribosomal biogenesis underlie age-related anabolic resistance to exercise in humans. The Journal of Physiology. 2016;594(24):7399–7417. doi:10.1113/JP272857

72. Edman S, Jones Iii RG, Jannig PR, et al. The 24-hour molecular landscape after exercise in humans reveals MYC is sufficient for muscle growth. EMBO Rep. 2024;25(12):5810–5837. doi:10.1038/s44319-024-00299-z

73. III RGJ, Edman S, Serrano N, et al. Making Sense of MYC in Skeletal Muscle: Location, Duration, and Magnitude. American Journal of Physiology-Cell Physiology. Published online June 27, 2025. doi:10.1152/ajpcell.00528.2025

74. Zhu X, Yeadon, James E., and Burden SJ. AML1 Is Expressed in Skeletal Muscle and Is Regulated by Innervation. Molecular and Cellular Biology. 1994;14(12):8051–8057. doi:10.1128/mcb.14.12.8051-8057.1994

75. Wang X, Blagden C, Fan J, et al. Runx1 prevents wasting, myofibrillar disorganization, and autophagy of skeletal muscle. Genes Dev. 2005;19(14):1715–1722. doi:10.1101/gad.1318305

76. Cai X, Gao L, Teng L, et al. Runx1 deficiency decreases ribosome biogenesis and confers stress resistance to hematopoietic stem and progenitor cells. Cell Stem Cell. 2015;17(2):165–177. doi:10.1016/j.stem.2015.06.002

77. Murach KA, Englund DA, Chambers TL, et al. A satellite cell-dependent epigenetic fingerprint in skeletal muscle identity genes after lifelong physical activity. The FASEB Journal. 2025;39(5):e70435. doi:10.1096/fj.202500177R

78. Anastasiadi D, Esteve-Codina A, Piferrer F. Consistent inverse correlation between DNA methylation of the first intron and gene expression across tissues and species. Epigenetics & Chromatin. 2018;11(1):37. doi:10.1186/s13072-018-0205-1

79. Straussman R, Nejman D, Roberts D, et al. Developmental programming of CpG island methylation profiles in the human genome. Nat Struct Mol Biol. 2009;16(5):564–571. doi:10.1038/nsmb.1594

80. Murach KA, Dimet-Wiley AL, Wen Y, et al. Late-life exercise mitigates skeletal muscle epigenetic aging. Aging Cell. 2022;21(1):e13527. doi:10.1111/acel.13527

81. Voisin S, Jacques M, Landen S, et al. Meta-analysis of genome-wide DNA methylation and integrative omics of age in human skeletal muscle. *Journal of Cachexia*, Sarcopenia and Muscle. 2021;12(4):1064–1078. doi:10.1002/jcsm.12741

82. Voisin S, Seale K, Jacques M, et al. Exercise is associated with younger methylome and transcriptome profiles in human skeletal muscle. Aging Cell. 2023;23(1):e13859. doi:10.1111/acel.13859

83. Turner DC, Gorski PP, Maasar MF, et al. DNA methylation across the genome in aged human skeletal muscle tissue and muscle-derived cells: the role of HOX genes and physical activity. Sci Rep. 2020;10(1):15360. doi:10.1038/s41598-020-72730-z

84. Wang S, Sun H, Ma J, et al. Target analysis by integration of transcriptome and ChIP-seq data with BETA. Nat Protoc. 2013;8(12):2502–2515. doi:10.1038/nprot.2013.150

85. Chambers TL, Wells J, Koopmans PJ, et al. At the Nexus Between Epigenetics and Senescence: The Effects of Senolytic (BI01) Administration on DNA Methylation Clock Age and the Methylome in Aged and Regenerated Skeletal Muscle. Aging Cell. 2025;24(7):e70068. doi:10.1111/acel.70068

86. Chambers TL, Dimet-Wiley A, Keeble AR, et al. Methylome–proteome integration after late-life voluntary exercise training reveals regulation and target information for improved skeletal muscle health. The Journal of Physiology. 2025;603(1):211–237. doi:10.1113/JP286681

87. Turner DC, Seaborne RA, Sharples AP. Comparative Transcriptome and Methylome Analysis in Human Skeletal Muscle Anabolism, Hypertrophy and Epigenetic Memory. Sci Rep. 2019;9(1):4251. doi:10.1038/s41598-019-40787-0

88. Voisin S, Jacques M, Landen S, et al. Meta-analysis of genome-wide DNA methylation and integrative omics of age in human skeletal muscle. J Cachexia Sarcopenia Muscle. 2021;12(4):1064–1078. doi:10.1002/jcsm.12741

89. Blocquiaux S, Ramaekers M, Van Thienen R, et al. Recurrent training rejuvenates and enhances transcriptome and methylome responses in young and older human muscle. JCSM Rapid Communications. 2022;5(1):10–32. doi:10.1002/rco2.52

90. Cheah MS, Wallace CD, Hoffman RM. Hypomethylation of DNA in human cancer cells: a site-specific change in the c-myc oncogene. J Natl Cancer Inst. 1984;73(5):1057–1065.

91. de Souza CRT, Leal MF, Calcagno DQ, et al. MYC deregulation in gastric cancer and its clinicopathological implications. PLoS One. 2013;8(5):e64420. doi:10.1371/journal.pone.0064420

92. Kaneko Y, Shibuya M, Nakayama T, et al. Hypomethylation of c-myc and epidermal growth factor receptor genes in human hepatocellular carcinoma and fetal liver. Jpn J Cancer Res. 1985;76(12):1136–1140.

93. Rao PM, Antony A, Rajalakshmi S, Sarma DS. Studies on hypomethylation of liver DNA during early stages of chemical carcinogenesis in rat liver. Carcinogenesis. 1989;10(5):933–937. doi:10.1093/carcin/10.5.933

94. Tsujiuchi T, Tsutsumi M, Sasaki Y, Takahama M, Konishi Y. Hypomethylation of CpG sites and c-myc gene overexpression in hepatocellular carcinomas, but not hyperplastic nodules, induced by a choline-deficient L-amino acid-defined diet in rats. Jpn J Cancer Res. 1999;90(9):909–913. doi:10.1111/j.1349-7006.1999.tb00834.x

95. Forsburg SL. Eukaryotic MCM Proteins: Beyond Replication Initiation. Microbiol Mol Biol Rev. 2004;68(1):109–131. doi:10.1128/MMBR.68.1.109-131.2004

96. Malysa A, Zhang XM, Bepler G. Minichromosome Maintenance Proteins: From DNA Replication to the DNA Damage Response. Cells. 2024;14(1):12. doi:10.3390/cells14010012

97. Borowik AK, Davidyan A, Peelor FF III, et al. Skeletal Muscle Nuclei in Mice are not Post-mitotic. Function. 2023;4(1):zqac059. doi:10.1093/function/zqac059

98. Conti A, Riva N, Pesca M, et al. Increased expression of Myosin binding protein H in the skeletal muscle of amyotrophic lateral sclerosis patients. Biochimica et Biophysica Acta (BBA) - Molecular Basis of Disease. 2014;1842(1):99–106. doi:10.1016/j.bbadis.2013.10.013

99. Saini SK, Pérez-Cremades D, Cheng HS, et al. Dysregulated Genes, MicroRNAs, Biological Pathways, and Gastrocnemius Muscle Fiber Types Associated With Progression of Peripheral Artery Disease: A Preliminary Analysis. Journal of the American Heart Association. 2022;11(21):e023085. doi:10.1161/JAHA.121.023085

100. Graber TG, Maroto R, Thompson JK, et al. Skeletal Muscle Transcriptome Alterations Related to Declining Physical Function in Older Mice. Journal of Ageing and Longevity. 2023;3(2):159–178. doi:10.3390/jal3020013

101. Chen J, Billings SE, Nishimune H. Calcium Channels Link the Muscle-Derived Synapse Organizer Laminin β2 to Bassoon and CAST/Erc2 to Organize Presynaptic Active Zones. J Neurosci. 2011;31(2):512–525. doi:10.1523/JNEUROSCI.3771-10.2011

102. Pillon NJ, Gabriel BM, Dollet L, et al. Transcriptomic profiling of skeletal muscle adaptations to exercise and inactivity. Nat Commun. 2020;11(1):470. doi:10.1038/s41467-019-13869-w

103. Smith JAB, Gabriel BM, Brady AJ, et al. Inactivity-induced NR4A3 downregulation in human skeletal muscle affects glucose metabolism and translation: Insights from *in vitro* analysis. Molecular Metabolism. 2025;99:102200. doi:10.1016/j.molmet.2025.102200

104. Sabaratnam R, Pedersen AJ, Eskildsen TV, Kristensen JM, Wojtaszewski JFP, Højlund K. Exercise Induction of Key Transcriptional Regulators of Metabolic Adaptation in Muscle Is Preserved in Type 2 Diabetes. J Clin Endocrinol Metab. 2019;104(10):4909–4920. doi:10.1210/jc.2018-02679

105. Katayama M, Nomura K, Mudry JM, Chibalin AV, Krook A, Zierath JR. Exercise-induced methylation of the *Serhl2* promoter and implication for lipid metabolism in rat skeletal muscle. Molecular Metabolism. 2025;92:102081. doi:10.1016/j.molmet.2024.102081

106. Pendergrast LA, Ashcroft SP, Ehrlich AM, et al. Metabolic plasticity and obesity-associated changes in diurnal postexercise metabolism in mice. Metabolism. 2024;155:155834. doi:10.1016/j.metabol.2024.155834

107. Petrany MJ, Swoboda CO, Sun C, et al. Single-nucleus RNA-seq identifies transcriptional heterogeneity in multinucleated skeletal myofibers. Nat Commun. 2020;11(1):6374. doi:10.1038/s41467-020-20063-w

108. Lagerwaard B, Nieuwenhuizen AG, Bunschoten A, de Boer VCJ, Keijer J. Matrisome, innervation and oxidative metabolism affected in older compared with younger males with similar physical activity. Journal of Cachexia, Sarcopenia and Muscle. 2021;12(5):1214–1231. doi:10.1002/jcsm.12753

109. Zhang Y, Santos MD, Huang H, et al. A molecular pathway for cancer cachexia-induced muscle atrophy revealed at single-nucleus resolution. Cell Reports. 2024;43(8). doi:10.1016/j.celrep.2024.114587

110. Lin H, Ma X, Sun Y, et al. Decoding the transcriptome of denervated muscle at single-nucleus resolution. J Cachexia Sarcopenia Muscle. 2022;13(4):2102–2117. doi:10.1002/jcsm.13023

111. Hepple RT, Rice CL. Innervation and neuromuscular control in ageing skeletal muscle. The Journal of Physiology. 2016;594(8):1965–1978. doi:10.1113/JP270561

112. Horwath O, Moberg M, Edman S, Philp A, Apró W. Ageing leads to selective type II myofibre deterioration and denervation independent of reinnervative capacity in human skeletal muscle. Experimental Physiology. 2025;110(2):277–292. doi:10.1113/EP092222

113. Sun C, Swoboda CO, Morales FM, et al. Lineage tracing of nuclei in skeletal myofibers uncovers distinct transcripts and interplay between myonuclear populations. Nat Commun. 2024;15(1):9372. doi:10.1038/s41467-024-53510-z

114. Murach KA, White SH, Wen Y, et al. Differential requirement for satellite cells during overload-induced muscle hypertrophy in growing versus mature mice. Skeletal Muscle. 2017;7(1):14. doi:10.1186/s13395-017-0132-z

115. Fry CS, Lee JD, Mula J, et al. Inducible depletion of satellite cells in adult, sedentary mice impairs muscle regenerative capacity but does not contribute to sarcopenia. Nat Med. 2015;21(1):76–80. doi:10.1038/nm.3710

116. Mylona E, Jones KA, Mills ST, Pavlath GK. CD44 regulates myoblast migration and differentiation. Journal of Cellular Physiology. 2006;209(2):314–321. doi:10.1002/jcp.20724

117. Roman W, Pinheiro H, Pimentel MR, et al. Muscle repair after physiological damage relies on nuclear migration for cellular reconstruction. Science. 2021;374(6565):355-359. doi:10.1126/science.abe5620

118. Carramusa L, Contino F, Ferro A, et al. The PVT-1 oncogene is a Myc protein target that is overexpressed in transformed cells. J Cell Physiol. 2007;213(2):511–518. doi:10.1002/jcp.21133

119. Soendenbroe C, Bechshøft CJL, Heisterberg MF, et al. Key Components of Human Myofibre Denervation and Neuromuscular Junction Stability are Modulated by Age and Exercise. Cells. 2020;9(4):893. doi:10.3390/cells9040893

120. Figueiredo VC, McCarthy JJ. Regulation of Ribosome Biogenesis in Skeletal Muscle Hypertrophy. Physiology (Bethesda). 2019;34(1):30–42. doi:10.1152/physiol.00034.2018

121. Soendenbroe C, Heisterberg MF, Schjerling P, et al. Molecular indicators of denervation in aging human skeletal muscle. Muscle & Nerve. 2019;60(4):453–463. doi:10.1002/mus.26638

122. Soendenbroe C, Andersen JL, Mackey AL. Muscle-nerve communication and the molecular assessment of human skeletal muscle denervation with aging. American Journal of Physiology-Cell Physiology. 2021;321(2):C317–C329. doi:10.1152/ajpcell.00174.2021

123. Sarto F, Franchi MV, McPhee JS, et al. Neuromuscular impairment at different stages of human sarcopenia. Journal of Cachexia, Sarcopenia and Muscle. 2024;15(5):1797–1810. doi:10.1002/jcsm.13531

124. McCarthy JJ, Srikuea R, Kirby TJ, Peterson CA, Esser KA. Inducible Cre transgenic mouse strain for skeletal muscle-specific gene targeting. Skelet Muscle. 2012;2:8. doi:10.1186/2044-5040-2-8

125. Stokes T, Cen HH, Kapranov P, et al. Transcriptomics for Clinical and Experimental Biology Research: Hang on a Seq. Advanced Genetics. 2023;4(2):2200024. doi:10.1002/ggn2.202200024

126. Jones III RG, Dimet-Wiley A, Haghani A, et al. A molecular signature defining exercise adaptation with ageing and in vivo partial reprogramming in skeletal muscle. The Journal of Physiology. 2023;601(4):763–782. doi:10.1113/JP283836

127. Tang Q, Chen Y, Meyer C, et al. A Comprehensive View of Nuclear Receptor Cancer Cistromes. Cancer Res. 2011;71(22):6940–6947. doi:10.1158/0008-5472.CAN-11-2091

128. Fleming SJ, Chaffin MD, Arduini A, et al. Unsupervised removal of systematic background noise from droplet-based single-cell experiments using CellBender. Nat Methods. 2023;20(9):1323–1335. doi:10.1038/s41592-023-01943-7

129. Young MD, Behjati S. SoupX removes ambient RNA contamination from droplet-based single-cell RNA sequencing data. GigaScience. 2020;9(12):giaa151. doi:10.1093/gigascience/giaa151

130. Hao Y, Stuart T, Kowalski MH, et al. Dictionary learning for integrative, multimodal and scalable single-cell analysis. Nat Biotechnol. 2024;42(2):293–304. doi:10.1038/s41587-023-01767-y

