## Supplementary Figures 1-4 for "The Age-Dependent Resident Myonuclear Multi-Omic Response to a Skeletal Muscle Hypertrophic Stimulus"

### Supplemental Figure 1

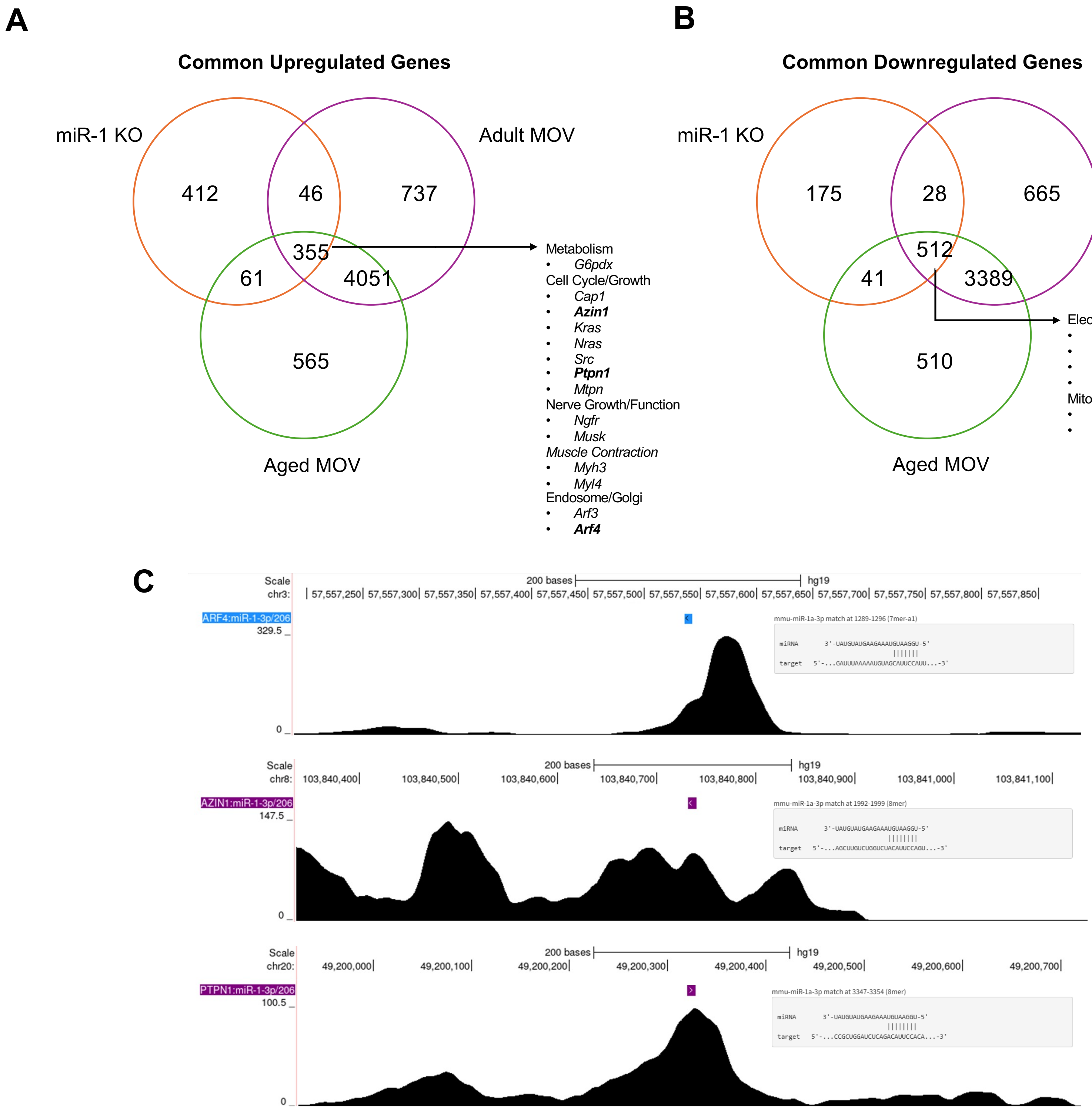

### Supplemental Figure 2

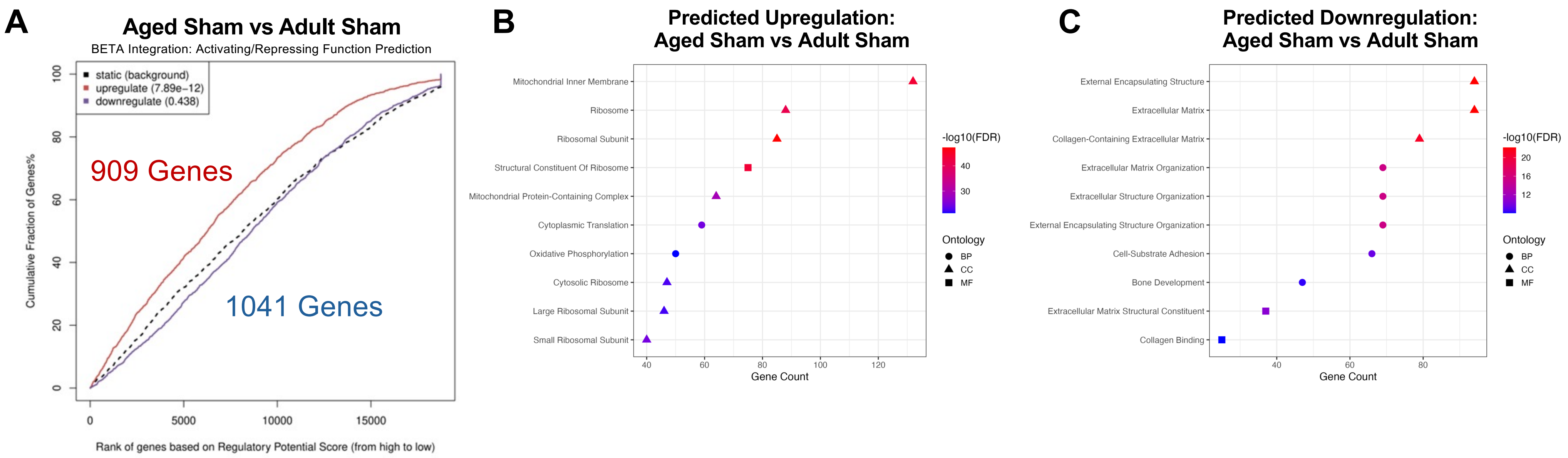

**Supplemental Figure 3**

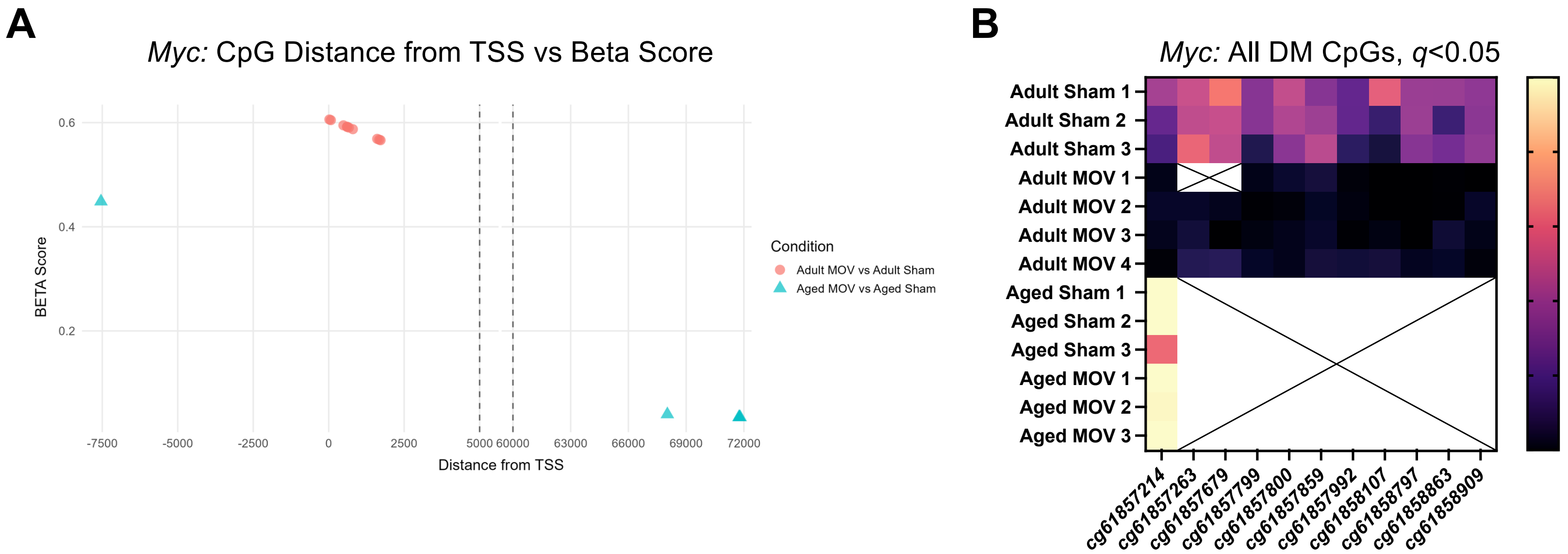


**C**

| Gene | Adult MOV: # BETA Reg. CpG Sites | Adult MOV: Avg. Dist. from TSS | Aged MOV: # BETA Reg. CpG Sites | Aged MOV: Avg. Dist. fr |
|---|---|---|---|---|
| *Ankrd1* | 11 | 48038 | 1 | 92080 |
| *Atf3* | 36 | 24567 | 7 | 53192 |
| *Runx1* | 19 | 11309 | 17 | 22361 |
| *Enah* | 21 | 7516 | 4 | 72183 |
| *Mybph* | 13 | 55646 | 0 | NA |
| *Myc* | 10 | 827 | 5 | 58176 |
| *Igf2bp2* | 22 | 3052 | 6 | 26020 |

#### Supplemental Figure 4

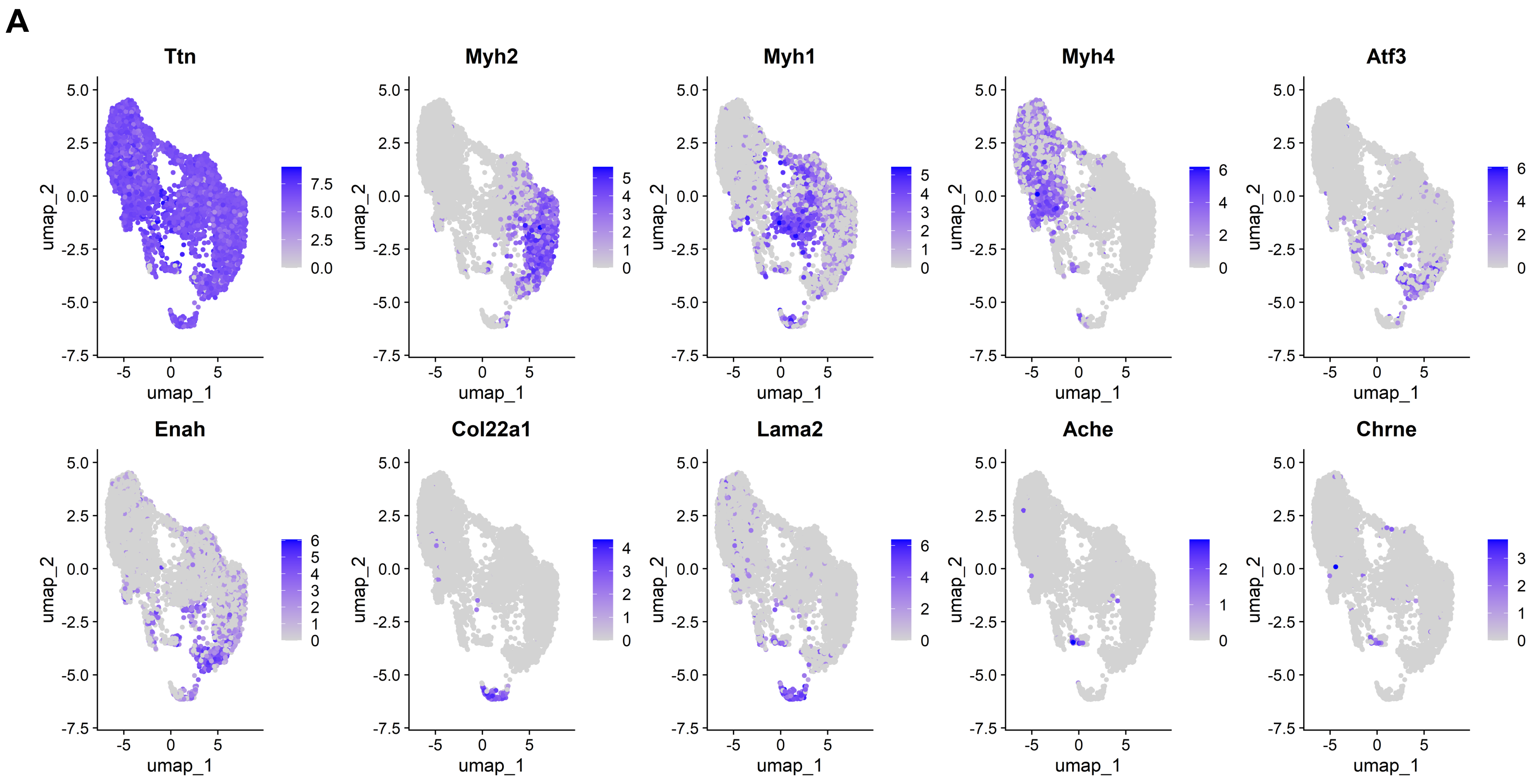
